# A library for fMRI real-time processing systems in python (RTPSpy) with comprehensive online noise reduction, fast and accurate anatomical image processing, and online processing simulation

**DOI:** 10.1101/2021.12.13.472468

**Authors:** Masaya Misaki, Jerzy Bodurka, Martin P Paulus

## Abstract

Real-time fMRI (rtfMRI)has enormous potential for both mechanistic brain imaging studies or treatment-oriented neuromodulation. However, the adaption of rtfMRI has been limited due to technical difficulties implement an efficient computational framework. Here, we introduce a python library for real-time fMRI (rtfMRI) data processing systems, Real-Time Processing System in python (RTPSpy), to provide building blocks for a custom rtfMRI application with extensive and advanced functionalities. RTPSpy is a library package including 1) a fast, comprehensive, and flexible online fMRI image processing modules comparable to offline denoising, 2) utilities for fast and accurate anatomical image processing to define an anatomical target region, 3) a simulation system of online fMRI processing to optimize a pipeline and target signal calculation, 4) simple interface to an external application for feedback presentation, and 5) a boilerplate graphical user interface (GUI) integrating operations with RTPSpy library. The fast and accurate anatomical image processing utility wraps external tools, including FastSurfer, ANTs, and AFNI, to make tissue segmentation and region of interest masks. We confirmed that the quality of the output masks was comparable with FreeSurfer, and the anatomical image processing could complete in a few minutes. The modular nature of RTPSpy provides the ability to use it for a simulation analysis to optimize a processing pipeline and target signal calculation. We present a sample script for building a real-time processing pipeline and running a simulation using RTPSpy. The library also offers a simple signal exchange mechanism with an external application using a TCP/IP socket. While the main components of the RTPSpy are the library modules, we also provide a GUI class for easy access to the RTPSpy functions. The boilerplate GUI application provided with the package allows users to develop a customized rtfMRI application with minimum scripting labor. The limitations of the package as it relates to environment-specific implementations are discussed. These library components can be customized and can be used in parts. Taken together, RTPSpy is an efficient and adaptable option for developing rtfMRI applications. The package is available from GitHub (https://github.com/mamisaki/RTPSpy) with GPL3 license.

## 1 Introduction

Online evaluation of human brain activity with real-time functional magnetic resonance imaging (rtfMRI) expands the possibility of neuroimaging. Its application has been extended from on-site quality assurance (1), brain-computer-interface (BCI) (2), brain self-regulation with neurofeedback (3), and online optimization in brain stimulation (4). Nevertheless, a complex system setup specific to an individual environment and noisy online evaluation of neural activation due to a limited realtime fMRI signal processing have hindered the utility of rtfMRI applications and reproducibility of its result (5). Indeed, the significant risk of noise contamination in the neurofeedback signal has been demonstrated in recent studies (6, 7). These issues have been addressed with a community effort releasing easy-to-use rtfMRI frameworks (1, 8–13) and consensus on reporting detailed online processing and experimental setups (14).

As one of the contributions to such an effort, we introduce a software library for rtfMRI; fMRI Real-Time Processing System in python (RTPSpy). The goal of the RTPSpy is to provide building blocks for making a highly customized and advanced rtfMRI system. The library is not assumed to provide a complete application package but offers rtfMRI data processing components to be used as a part of a user’s custom application. We suppose that the tools of RTPSpy can also be combined with other frameworks as a part of processing modules.

RTPSpy is a python library that includes a fast and comprehensive online fMRI image processing pipeline comparable to offline processing (7) and an interface module for an external application to receive real-time brain activation signals via TCP/IP socket. Each online data processing component is implemented in an independent class, and a processing pipeline can be created by chaining these modules. In addition to the online fMRI signal processing modules, the library provides several utility modules, including brain anatomical image processing tools for fast and accurate tissue segmentation, and an online fMRI processing simulation system. Although these utilities may not always be required in a rtfMRI session, the fast anatomical image processing can be useful for identifying anatomically defined target regions, and the simulation analysis is vital for building an optimal processing pipeline (7, 15, 16). We also provide a boilerplate graphical user interface (GUI) application integrating operations with RTPSpy, and a sample application of neurofeedback presentation using PsychoPy (17) to demonstrate how the RTPSpy is implemented in an application and to interface to another external application. The GUI application is presented as just one example of library usage. However, a user may develop a custom neurofeedback application with minimum modification on this example script.

The aim of this paper is to introduce the structure of the RTPSpy library and its usages as a part of a neurofeedback application. We hope that RTPSpy is used as a part of a user’s own custom application so that the current report focuses on how to script the RTPSpy online processing pipeline and implement it in an application. The detailed usage of the example application is presented in GitHub (https://github.com/mamisaki/RTPSpy/tree/main/example/ROI-NF). Also, this paper does not provide a comprehensive evaluation of the library’s performance in detail. Such evaluations have been done in our previous report (7), and only a short overview of the previous report was given in this report. Comparison with other exiting rtfMRI frameworks is also out of the scope of this paper. We recognize that many excellent packages are released for rtfMRI (1, 8–13), and we do not claim that RTPSpy is the best. The claim of the advanced functionality of the RTPSpy is for its own sake and not relative to other tools. RTPSpy and this paper aim to offer users another option for developing a custom rtfMRI application.

This paper is organized as follows. The next section summarizes the installation and supporting system information. The third section introduces the online fMRI data processing modules in RTPSpy, the main components of the library. The issues and caveats in online fMRI data analysis and how they are addressed in RTPspy implementation are discussed here. The fourth section describes fast and accurate anatomical image processing tools. A custom processing stream was made by wrapping external tools, FastSurfer (18), AFNI (https://afni.nimh.nih.gov/), and ANTs (http://stnava.github.io/ANTs/). We also evaluated the accuracy of tissue segmentation and the quality of tissue-based noise regressors made by this stream compared to FreeSurfer segmentation. The fifth and sixth sections illustrate the usage of library classes to build a processing pipeline and run a simulation analysis. An example GUI implementation is presented in the seventh section. The last section discusses the system components that are not provided with RTPSpy but are required for a complete system depending on an individual environment. The RTPSpy can be obtained from GitHub (https://github.com/mamisaki/RTPSpy) with GPL3 license.

## 2 Installation and supporting systems

RTPSpy is assumed to be run on a miniconda3 (https://docs.conda.io) or Anaconda (https://www.anaconda.com/) environment. A yaml file for installing the required python libraries in an anaconda environment is provided with the package for easy installation. RTPSpy’s anatomical image processing depends on several external tools, AFNI (https://afni.nimh.nih.gov/), FastSurfer (https://deep-mi.org/research/fastsurfer/), and ANTs (https://pypi.org/project/antspyx/). While AFNI needs to be installed separately, FastSurfer and ANTs installation is integrated into the RTPSpy setup. Refer to the GitHub site (https://github.com/mamisaki/RTPSpy) for further details.

RTPSpy can take advantage of graphical processing unit (GPU) computation. GPU can be utilized in the online fMRI data processing and anatomical image processing with FastSurfer (https://deep-mi.org/research/fastsurfer/). To use GPU computation, a user needs a GPU compatible with NVIDIA’s CUDA toolkit (https://developer.nvidia.com/cuda-toolkit) and to install a GPU driver compatible with the CUDA. The CUDA toolkit will be installed with the yaml file. We note that GPU is not mandatory for RTPSpy. Online data processing speed in RTPSpy is fast enough for realtime fMRI even without GPU, while GPU can enhance it further (Section 3.2). Also, RTPSpy provides an alternative anatomical image processing stream not using FastSurfer, while the image segmentation accuracy is better with FastSeg (Section 4.2).

RTPSpy has been developed on a Linux system (Ubuntu 20.04). It can also be run on Mac OS X and Windows with the Windows Subsystem for Linux (WSL), while GPU computation is not supported on OS X and WSL for now.

## 3 RTPSpy online fMRI data processing

### 3.1 Overview of the library design

Figure 1 shows an overview of the modules composing an online processing pipeline. RTPSpy includes six online fMRI data processing modules; a real-time data loader (RTP_WATCH), slicetiming correction (RTP_TSHIFT), motion correction (RTP_VOLREG), spatial smoothing (RTP_SMOOTH), noise regression (RTP_REGRESS), and an application module (RTP_APP). A utility module for an external application to receive a processed signal, RTP_SERV, is also provided. RTP_WATCH is the entrance module, and RTP_APP is the terminal module of a pipeline. Other modules have common input and output interfaces so that they can be connected in any combination and order. For example, when a conventional pipeline only with a motion correction is enough, the pipeline can be made only with RTP_WATCH, RTP_VOLREG, and RTP_APP modules. If more comprehensive processing is required, all the components can be chained in a pipeline.

**Figure 1.**
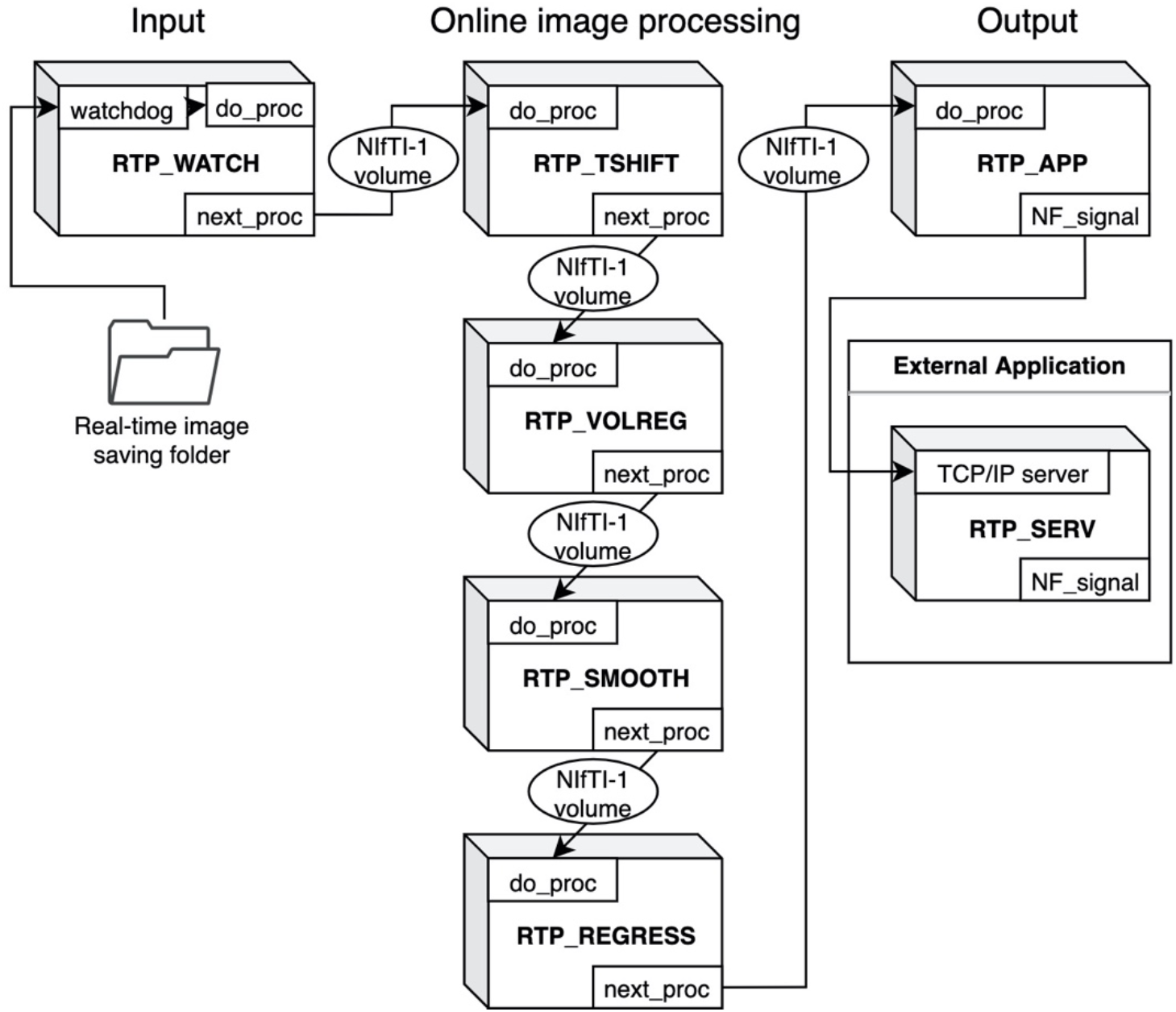
Overview of the RTPSpy module design for creating an online fMRI processing pipeline. RTP_WATCH is the entrance module, and RTP_APP is the terminal module of a pipeline. Other modules have common input and output interfaces. They can be connected in any combination and order. The modules exchange data with the Nifti1Image object of the whole-brain volume data. RTP_SERV is a utility module for an external application to receive a processed signal.

These modules are implemented as a python class. The module’s interface method is ‘do_proc’, which receives a NiBabel (https://nipy.org/nibabel/) Nifti1Image object. Its calling format is the same for all modules. The modules exchange data with the Nifti1Image object of the whole-brain volume data. The processing chain can be made by setting the ‘next_proc’ property to an object of the next module. Calling the ‘do_proc’ method at the head of the pipeline calls the next module’s ‘do_proc’ method in the chain. This simple function interface enables the easy creation of a custom pipeline (see Section 5.1 for details).

We assume that the input and output parts should be customized according to the user’s environment and an application need. For example, if a user wants to use another real-time image feeding (e.g., a dicom export feature of a scanner), RTP_WATCH can be replaced or modified in a preferred way. Also, the RTP_APP can be customized to calculate a neurofeedback signal in a user’s way. An example script for such customization is presented in Section 7. We note that RTPSpy is not intended to provide a complete application for any environment. Instead, a necessary module for each environment is supposed to be developed by a user. RTPSpy offers a framework of the interface and building blocks of online fMRI data processing.

### 3.2 Real-time performance

Retaining the whole-brain data throughout the pipeline enables a common interface between modules. It also provides freedom of neurofeedback signal calculation (i.e., an ROI average, connectivity of multiple regions, and multi-voxel patterns in the whole brain) with various combinations of processing modules. Although this implementation seemed burdensome for realtime computation, we found that processing whole-brain volume does not significantly affect the real-time performance in RTPSpy. Our previous report (7) showed that the pipeline processing was completed in less than 400 ms on a current PC equipped with a GPU. Here, we also evaluated the processing time with several PC specifications with and without GPU for a sample fMRI data (128 × 128 × 34 matrix, 203 volumes). Note that this evaluation is not a comprehensive performance test but rather a rough guide to the PC specifications required for real-time processing with RTPSpy.

Table 1 shows the specifications of the tested PCs. ‘Linux+GPU’ is the same one used in Misaki and Bodurka (7). The evaluated pipeline included all modules implemented in RTPSpy and RTP_REGRESS includes all available regressors. Figure 1 shows processing times for each module. The figure shows the results after TR = 45 since the regression waited to receive 40 volumes, excluding the initial three volumes, and the processing of the first regressed volume took a long time due to initialization and retrospective processing (see Section 3.3, RTP_REGRESS). The processing time of RTP_WATCH is from a file creation time to send the volume data to the next module. The processing time of RTP_APP is to extract an ROI average signal and write it in a text file.

**Table 1.** PC specifications used for the computation time evaluation.

| Name | CPU | RAM | Storage | GPU |
| --- | --- | --- | --- | --- |
| Linux+GPU | Dual Intel Xeon Gold 6126, 2.6 GHz, 12-core | 256 GB, DDR4 | HDD | NVIDIA TITAN V |
| Linux | Dual Intel Xeon Gold 6126, 2.6 GHz, 12-core | 256 GB, DDR4 | HDD | No |
| MacBookPro1 | Intel Core i9, 2.3 GHz, 8-Core | 32 GB, DDR4 | SSD | No |
| MacBookPro2 | Intel Core i7, 2.7 GHz, 4-core | 16 GB, DDR3 | SSD | No |
| Windows1 | Intel Xeon W-2245, 3.9 GHz, 8-core | 32 GB, DDR4 | SSD | No |
| Windows2 | Intel Core i5, 2.0 GHz, 4-core | 8 GB, DDR3 | HDD | No |

The results were consistent with the previous report (7). The most time-consuming processing was RTP_VOLREG. RTP_REGRESS’s processing time increased with TR since cumulative GLM uses more samples in later TRs (see Section 3.3, RTP_REGRESS). The slope of the increase was the lowest with GPU, indicating that GPU can be beneficial when a scan has many volumes. Interestingly, however, the total processing time was not significantly different by GPU usage, and MacBookPro showed comparable performance with a high-end Linux PC, at least for the present scan length. The Windows showed relatively longer processing times regardless of the specification, which might be due to the overhead of the Windows subsystem for Linux. These results indicate that the PC requirement for RTPSpy is not high, at least for an ordinary real-time fMRI scan with a few seconds TR and less than a few hundred volumes.

Even if computation time does not limit real-time fMRI processing, the limited number of sample points available online poses a challenge for online processing yet. The next section describes the details of each module functionalities and online analysis methods in RTPSpy to address this issue.

### 3.3 Implementations of the online fMRI processing algorithms

Table 2 summarizes the functions of RTPSpy processing modules. The class files for these modules can be found in the ‘rtpspy’ directory of the package. The issue of the limited number of online available sample points is critical for slice timing correction, signal scaling, and online noise regression. This section describes the methods used in the RTPSpy modules to address this issue.

**Table 2.**
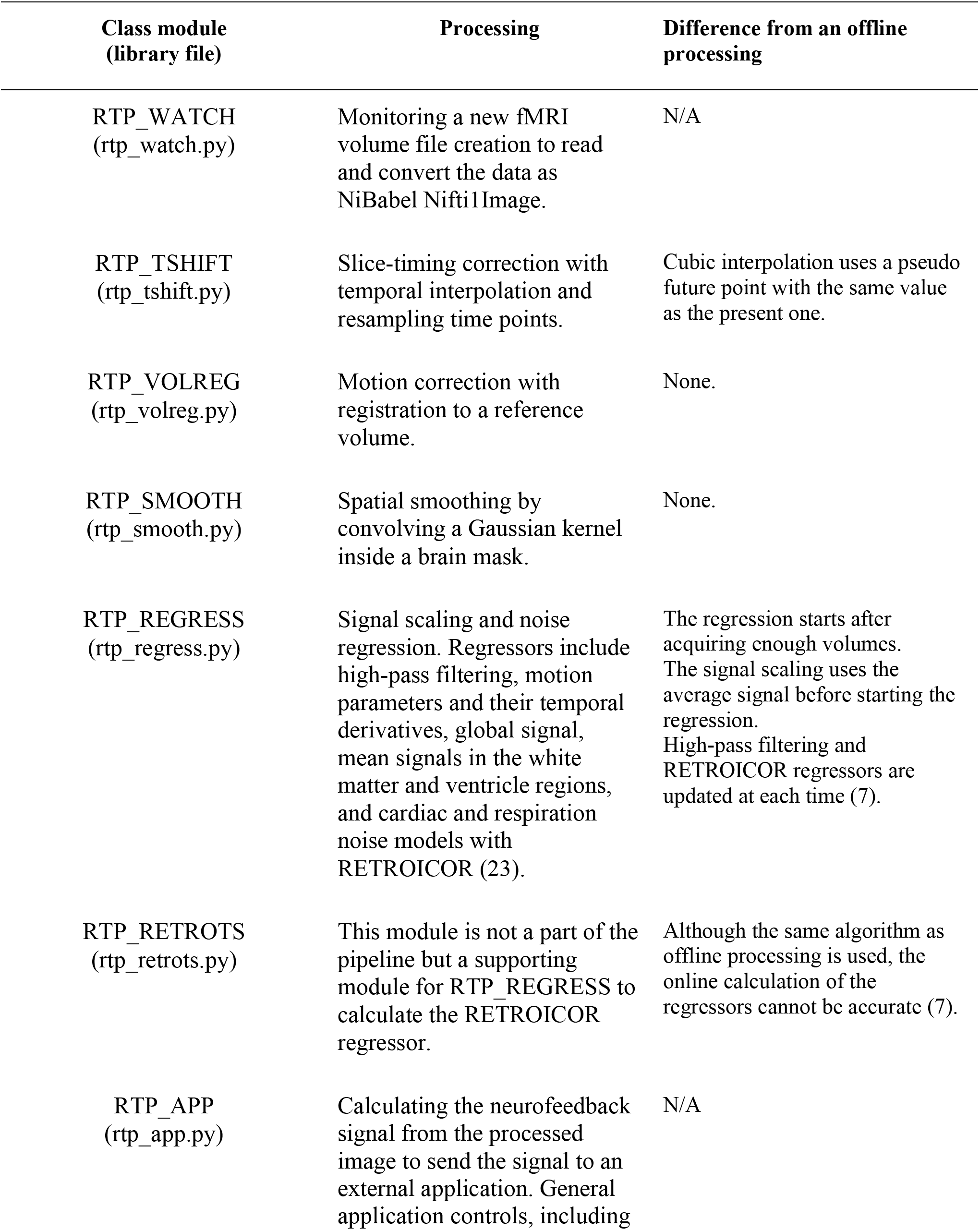
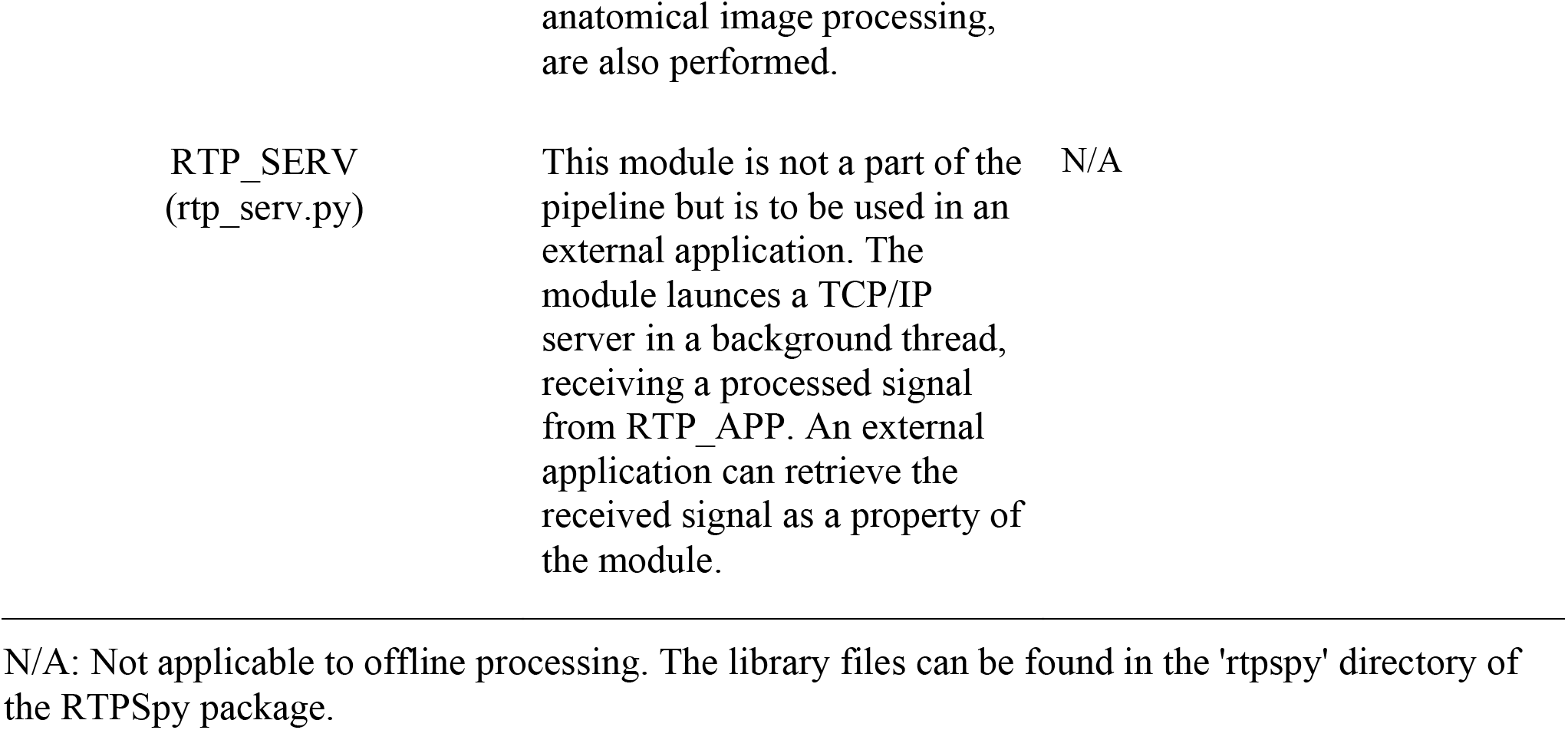
Summaries of RTPSpy real-time processing modules and their differences from offline processing.

**RTP_WATCH** finds a newly created file in a watching directory in real-time, reads the data, and sends it to the next processing module. The watchdog module in python (https://pythonhosted.org/watchdog/index.html) is used to detect a new file creation. RTP_WATCH uses the NiBabel library (https://nipy.org/nibabel/) to read the file. The currently supported file types are NIfTI, AFNI’s BRIK, and Siemens’ mosaic dicom. Technically, this module can handle all file types supported by NiBabel, so a user can easily extend the supported files as needed. The wholebrain volume data is retained in the NiBabel Nifti1Image format throughout the RTPSpy pipeline. The observer function used in RTP_WATCH (PollingObserverVFS) is system-call independent and can work with various file systems. However, polling may take a long time if many files are in the monitored directory, hindering real-time performance. If a user finds a significant delay by saving files of many runs in a single directory, it is recommended to clean or move files at each run.

Although RTP_WATCH offers a simple interface to read a data file in real-time, how the MRI scanner saves the reconstructed image varies across the manufacturers and sites. RTPSpy does not provide a universal solution for that. A user may need another package to send data to the watched directory or modify the script file (rtp_watch.py) to adjust for each environment. We will discuss this limitation of the environment-specific issues in the last section.

**RTP_TSHIFT** performs a slice-timing correction by aligning the signal sampling times in different slices to the same one with temporal interpolation. Because we cannot access the future time point in real-time, the online processing cannot be equivalent to offline. RTPSpy aligns the sampling time to the earliest slice for avoiding extrapolation. RTPSpy implements two interpolation methods, linear and cubic. The linear interpolation uses only the current and one past time point so that it is equivalent to offline processing. The cubic interpolation uses four time-points; two from the past, the present, and one future. RTPSpy puts a pseudo future point with the same value as the present one to perform the cubic interpolation. We have confirmed that this pseudo cubic method has a higher correlation with a high-order interpolation method (e.g., FFT) than a linear method (19). By default, RTPSpy uses cubic interpolation.

Slice-timing correction is often skipped in real-time fMRI processing, and its effect could be minor when TR is short (20, 21). However, some neurofeedback signal extraction methods, such as the two-point connectivity (22), could be sensitive to a small timing difference between slices. The two-point method evaluates the consistency of the signal change direction (increase/decrees) at each TR, which could be sensitive to the timing of signal direction change between ROIs in different slices. The user can specify the particular module of RTPSpy, which does not enforce the specific pipeline for the real-time fMRI processing. We also note that slice-timing correction takes no significant cost of computational time (Fig. 2).

**Figure 2.**
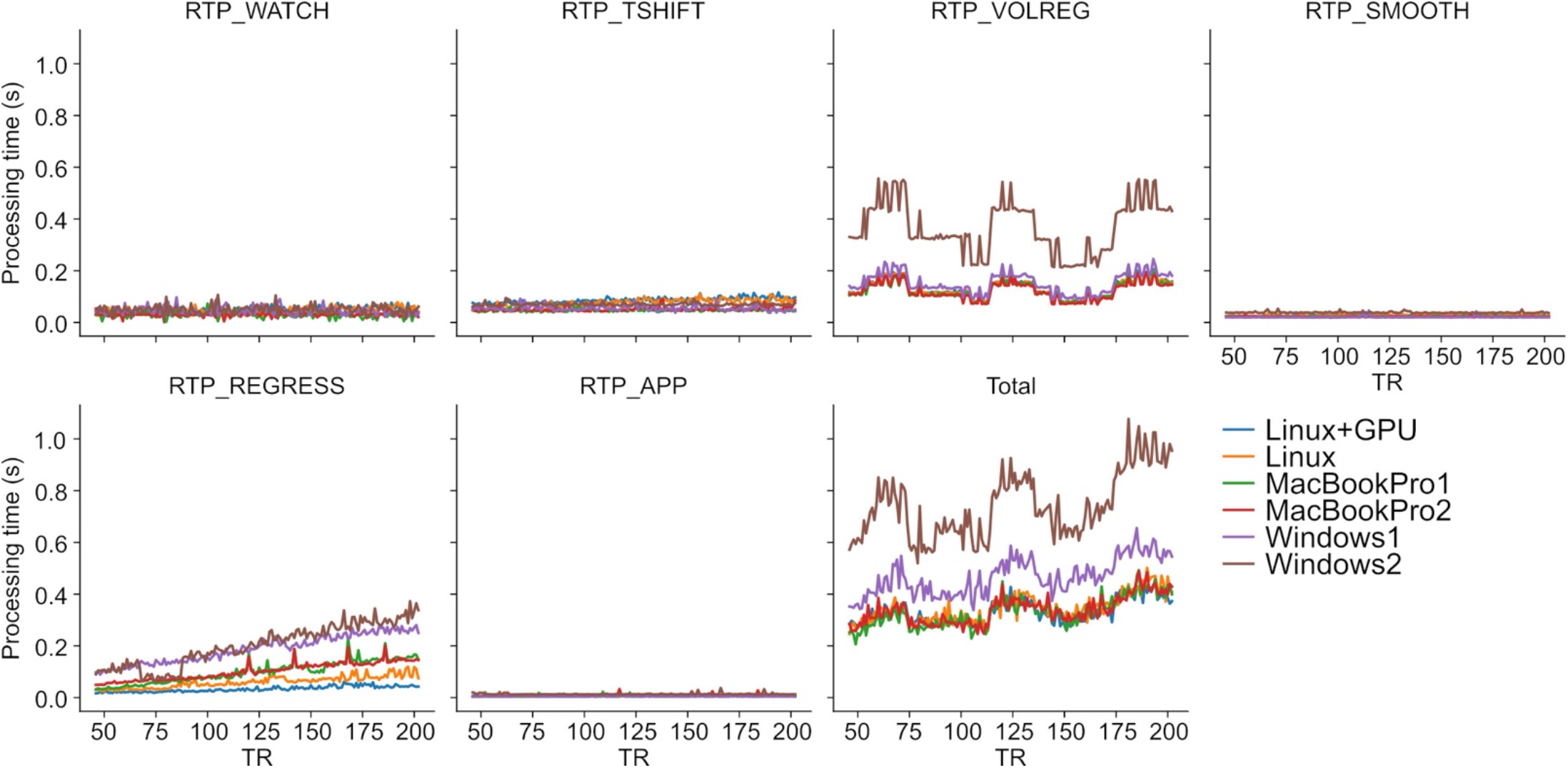
RTPSpy online fMRI data processing times. See Table 1 for the specification of the PCs. The evaluation was done with a sample fMRI data (128 × 128 × 34 matrix, 203 volumes). Processing with RTP_REGRESS included all available regressors (Legendre polynomials for high-pass filtering, 12 motion parameters, global signal, mean white matter and ventricle signals, and RETROICOR regressors).

**RTP_VOLREG** performs motion correction by registering volumes to a reference one. The same functions as the AFNI’s 3dvolreg (https://afni.nimh.nih.gov/pub/dist/doc/program_help/3dvolreg.html), a motion correction command in the AFNI toolkit, is implemented in RTP_VOLREG. We compiled the C source codes of 3dvolreg functions into a custom C shared library file (librtp.so in the RTPSpy package), and RTP_VOLREG accesses it via the python ctypes interface. Thus, this online processing is equivalent to the offline 3dvolreg. By default, RTP_VOLREG uses heptic (7th order polynomial) interpolation at image reslicing, the same as the 3dvolreg default.

**RTP_SMOOTH** performs spatial smoothing by convolving a Gaussian kernel within a masked region. Like RTP_VOLREG, RTPSpy uses the AFNI’s 3dBlurInMask (https://afni.nimh.nih.gov/pub/dist/doc/program_help/3dBlurInMask.html) functions compiled into a C shared library file (librtp.so), and accessed via ctypes interface in python. This process has no difference between online and offline processing.

**RTP_REGRESS** performs a signal scaling and noise regression analysis. The regression requires at least as many data points as regressors in the model and will not commence the process until sufficient number of data points have been collected. The signal scaling is done with the average signal in this waiting period and converts a signal into percent change relative to the average in each voxel. We note that this scaling is not equivalent to the offline processing using an average of all time points in a run so that the absolute amplitude cannot be comparable between the online and offline processing. We also note that the volumes before the start of regression are processed retrospectively so that the saved data includes all volumes. Once enough volumes are received, the regression is done with an ordinary least square (OLS) approach using the PyTorch library (https://pytorch.org/), which allows a seamless switching of CPU and GPU usage according to the system equipment. The residual of regression is obtained as a denoised signal.

The regressors can include high-pass filtering (Legendre polynomials), motion parameters (three shifts and three rotations), their temporal derivatives, global signal, mean signals in the white matter and ventricle regions, and cardiac and respiration noise models with RETROICOR (23). The order of the Legendre polynomials for high-pass filtering is adjusted according to the data length at each volume with 1 + int(d/150), where d is the scan duration in seconds (the default in AFNI). The motion parameters were received from the RTP_VOLREG module in real-time. The global signal and the mean white matter and ventricle signals are calculated from the unsmoothed data, which is also received from the RTP_VOLREG module. These regressors were made from the mask files defined in ‘GS_mask’ (global signal mask), ‘WM_mask’, and ‘Vent_mask’ properties of the module. As the RTP_REGRESS depends on RTP_VOLREG outputs, RTP_VOLREG must be included before RTP_REGRESS when it is used in a pipeline. A user can also include any pre-defined timeseries such as a task design as a covariate in the regressors. It is up to a user to decide which regressor to use. The report in Misaki and Bodurka (7) has shown which regressor was effective in reducing what noise in what brain regions and connectivity, which may help decide the noise regressor choice.

RTPSpy uses cumulative GLM (cGLM), which performs regression with all samples at each time, rather than incremental GLM (iGLM), which updates only the most recent estimates based on previous estimates (24). In Misaki and Bodurka (7), we indicated that high-pass filtering regressor, either Legendre polynomial or discrete cosine transform, filtered higher frequencies than the designed threshold at early TRs unless the regressor was adjusted at each TR. This adjustment requires a retrospective update of the regressor. Similarly, the online creation of RETROICOR regressors, made from real-time cardiac and respiration signal recordings, could not be accurate compared to the offline creation, and the error was accumulated unless retrospective correction was made (see Figs. 2 and 3 in Misaki and Bodurka (7)). RTPSpy uses cGLM because cGLM has the advantage of being able to recalculate regressors at each volume, thereby improving the quality of regressors made online with limited samples. Although this implementation, whole-brain processing with cGLM, seemed burdensome for real-time processing, the computation time is not inhibitive to real-time performance, as shown in Section 3.2.

**Figure 3.**
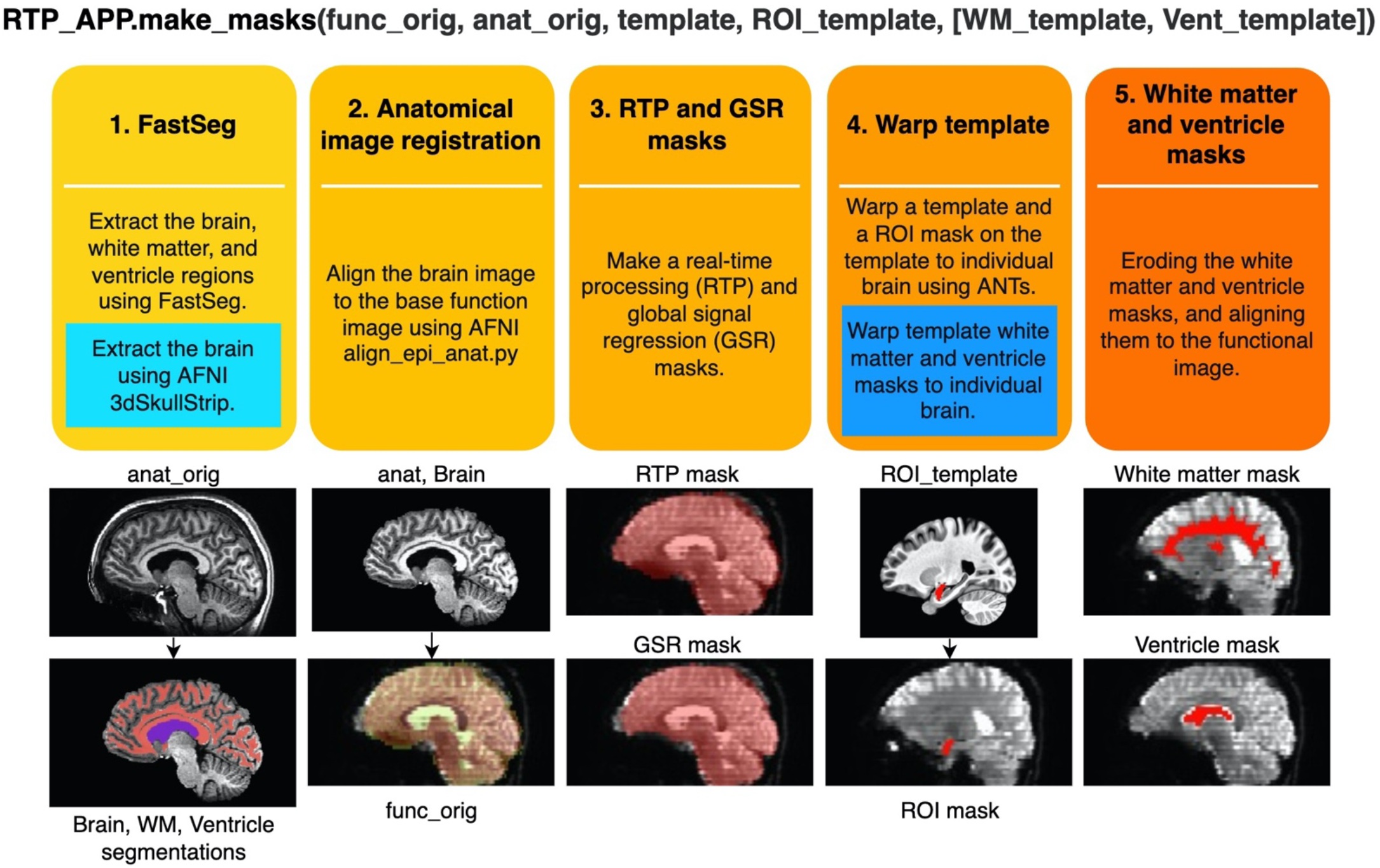
Procedures of creating image masks in the ‘make_masks’ method of RTP_APP. Two blue boxes show the procedures used in the alternative stream without using FastSeg. In the alternative stream, the first process is replaced by a blue box, and the fourth process is performed by adding the procedure in the blue box. The images below demonstrate what image processing is done at each step.

**RTP_RETROTS** is not a pipeline component (thus, not shown in Figure 1) but a supporting module for RTP_REGRESS to calculate the RETROICOR regressors from cardiac and respiration signals in real-time. In our environment, cardiac and respiration signals are measured using photoplethysmography and a pneumatic respiration belt, respectively. Although this hardware implementation could depend on the environment of each site (see Section 8), once the signal acquisition is set up, the usage of RTP_RETROTS is simple. Its interface method (‘do_proc’) receives respiration and cardiac signal values as one-dimensional arrays, the signals’ sampling frequency, and fMRI TR parameters. Then, the method returns the RETROICORE regressors.

RTP_RETROTS implements the same functions as the AFNI’s RetroTS.py script, which are rewritten in C codes and compiled into a shared library (librtp.so). The module makes four cardiac and four respiration noise basis regressors. It is possible to also create a respiration volume per time (RVT) regressors (25), however, we do not recommend using them in online processing. Our previous study (7) indicated that the online evaluation of RVT regressors could not be accurate, and its usage could introduce an artifactual signal fluctuation in the processed signal in an online regression.

**RTP_APP** receives the processed image and calculates the neurofeedback signal from it. The default implementation extracts the average signal in an ROI mask, defined in the ‘do_proc’ method of the rtp_app.py file. This method is provided as a prototype and can be customized according to the need for individual applications. Section 7 and Figure 10 show an example of a customized method. The RTPSpy noise reduction is performed for the whole-brain voxels, which is advantageous in calculating the feedback signals from multiple regions, such as the functional connectivity and decoding neurofeedback (26). The calculated signal can be sent to an external application through a network socket to the RTP_SERV module (Fig. 1; see also Section 7 and Figure 11). The RTP_APP class also implements general application control methods, including anatomical image processing described in the next section and a high-level scripting interface explained in Section 5.2.

**RTP_SERV** is not a part of the image processing pipeline but offers an interface class for an external application to communicate to RTPSpy. This module is assumed to be implemented in an external application as a receiver of the processed signal. Instantiating this class launches a TCP/IP server in a background thread in an external application to receive a real-time neurofeedback signal (see Section 7).

## 4 Anatomical image processing with fast and accurate tissue segmentation

Anatomical image processing is often required in a rtfMRI session. While there are several ways to define the target region for neurofeedback (e.g., functional localizer), if the target brain region is defined in the template brain with a group analysis, we need to warp the region mask into the participant’s brain. The noise regressions with the global signal and white matter and ventricle mean signals also require a brain mask and tissue segmentation masks on an individual brain image.

Although there are many tools for brain tissue segmentation using a signal intensity, they are prone to an image bias field and often need a manual correction. Another approach for tissue segmentation uses anatomical information to segment the regions in addition to the signal intensity, such as FreeSurfer (https://freesurfer.net/). FreeSurfer usually offers more accurate and robust segmentation than using only the signal intensity, but its process takes hours or longer to complete, inhibiting its use in a single visit rtfMRI session. Recently, an alternative approach of brain image segmentation using a deep neural network has been released as FastSurfer (18). FastSurfer uses a U-net architecture (27) trained to output a segmentation map equivalent to the FreeSurfer’s volume segmentation from an input of anatomical MRI image. FastSurfer can complete the segmentation in a few minutes with GPU. We made a script called FastSeg utilizing the advantage of FastSurfer to extract a brain mask (skull stripping), gray matter, white matter, and ventricle segmentation. FastSeg is implemented as part of the RTPSpy anatomical image processing pipeline and also released as an independent tool (https://github.com/mamisaki/FastSeg). The FastSurfer process in the FastSeg could take very long (about an hour) if GPU was not available. Therefore, RTPSpy also offers another processing stream that does not use FastSeg. This section describes the flow of the anatomical image processing steps and shows the evaluation results of their segmentation accuracy and noise regressor quality compared to FreeSurfer’s segmentation.

### 4.1 Anatomical image processing pipeline

RTPSpy offers a simple function interface to run a series of anatomical image processing, the ‘make_masks’ method in RTP_APP class. Figure 3 shows the processing pipeline in this method. The method receives filenames of a reference function image (func_orig), anatomical image (anat_orig), template image (template, optional), and a region of interest (ROI) image in the template space (ROI_template, optional). If the alternative processing stream without FastSeg is used, white matter and ventricle masks defined in the template space (WM_template, Vent_template) can also be received. The process includes the following five steps.

1. Extracting the brain (skull stripping), white matter, and ventricle regions using FastSeg. FastSeg uses the first stage of the FastSurfer process to make a volume segmentation map (DKTatlas+aseg.mgz). Then, all the segmented voxels are extracted as the brain mask with filling holes. The white matter mask is made with a union of the white matter and corpus callosum segmentations. The ventricle mask is made with lateral ventricle segmentation. We did not include small ventricle areas because the mask is used only for making a regressor for online fMRI denoising. In the alternative stream not using FastSeg (blue box in Fig. 3), AFNI’s 3dSkullStrip is used for brain extraction. White matter and ventricle masks are made in a later step (step 4) by warping the template masks into an individual brain.
2. Aligning the extracted brain image to a reference function image using AFNI align_epi_anat.py.
3. Aligning and resampling the brain mask into the reference function image space using the parameters made in step 2 and making a signal mask of the function image using 3dAutomask in AFNI. The union of these masks is made as a real-time processing mask (RTPmask.nii.gz), used at spatial smoothing in RTP_SMOOTH and defining the processing voxels in RTP_REGRESS. The intersect of these masks is also made as a global signal mask (GSRmask.nii.gz), used in RTP_REGRES.
4. If the template image and the ROI mask on the template are provided, the template brain is warped into the participant’s anatomical brain image using the python interface of ANTs registration (https://github.com/ANTsX/ANTsPy). Then, the ROI mask on the template is warped into the participant’s brain anatomy image and resampled to the reference function image to make an ROI mask in the functional image space. This mask will be used for neurofeedback signal calculation. In the alternative stream not using FastSeg (blue box in Fig. 3), white matter and ventricle masks defined in the template brain are also warped into an individual brain.
5. Eroding the white matter (two voxels) and ventricle (one voxel) masks and aligning them to the functional image space using the alignment parameters (affine transformation) estimated at step 2. These masks will be used for the white matter and ventricle average signal regression. These anatomical image processing could be completed in less than a few minutes. Table 3 shows the processing times with and without FastSeg on the PCs listed in Table 1 for one sample image (MPRAGE image with 256 × 256 × 120 matrix and 0.9 × 0.9 × 1.2 mm voxel size).

**Table 3.** Anatomical image processing times on the PCs shown in Table 1 (main text).

| PC | Linux+GPU* | Linux | MacBookPro1 | MacBookPro2 | Windows1 | Windows2 |
| --- | --- | --- | --- | --- | --- | --- |
| Processing time (s) | 89 | 146 | 197 | 188 | 241 | 473 |

\*: FastSeg was used in Linux+GPU. The alternative stream was used for others.

### 4.2 Evaluations for the segmentation masks and noise regressors

Since the anatomical segmentation by FastSurfer is not exactly the same as FreeSurfer (18), we evaluated the quality of white matter and ventricle masks made by the FatSeg compared to the FreeSurfer segmentation. The comparison was made for 87 healthy participants’ anatomical and resting-state fMRI images (age 18-55 years, 45 females) used in our previous study (7). We also performed the same comparison for the masks made by the alternative processing stream without FastSeg.

Figure 4 upper panel shows the Dice coefficients of the segmentation masks with the FreeSufer segmentation in anatomical image resolution. The masks made by FastSeg had high Dice coefficients with FreeSurfer segmentation showing their good agreement, while the masks made by the alternative stream had lower agreements, especially for the white matter mask. Nevertheless, the effect of these discrepancies could be minor in creating a noise regressor at functional image resolution. The bottom panel of Figure 4 shows the correlation between the average white matter and ventricle fMRI signals created from the FastSeg (or alternative stream) and FreeSurfer masks. For the FastSeg, the correlation was nearly 1.0 (higher than 0.98 even for the minimum sample). Although a few samples had a relatively low Dice coefficient for the FastSeg ventricle mask, that was because their ventricle region was small, and minor error affected the Dice coefficient much. Indeed, the signal correlation for the sample with minimum Dice coefficient (0.77) was as high as 0.99, indicating a minor segmentation error. Thus, the effect of the segmentation difference between the FastSeg and FreeSurfer was minor on the mean white matter and ventricle signals. The noise regressors made from the FastSeg hold equal quality to those made from FreeSurfer segmentations.

**Figure 4.**
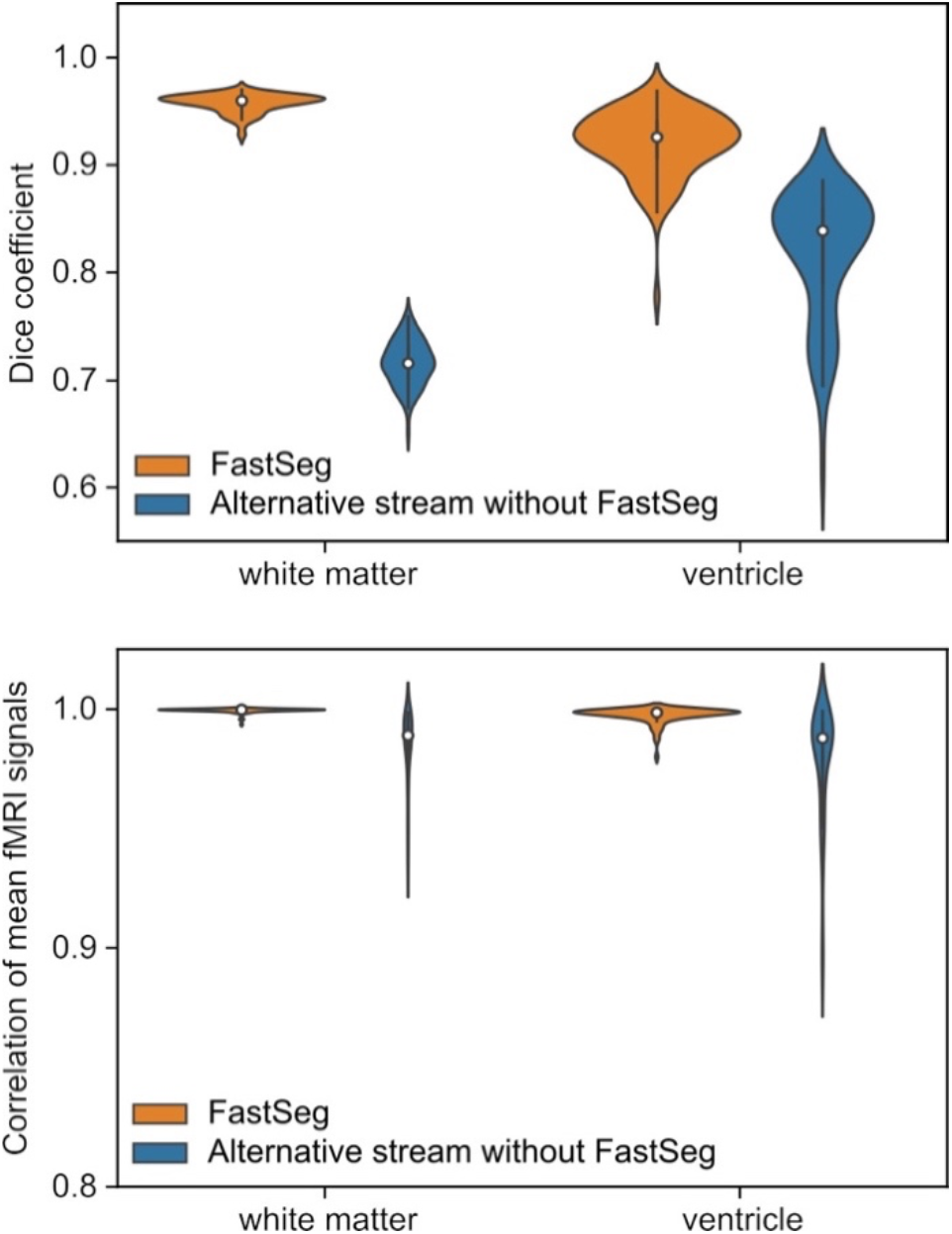
Quality evaluations for the masks created by the RTPSpy anatomical image processing streams by comparing to FreeSurfer segmentation. The upper panel shows Dice coefficients with FreeSurfer segmentations for the white matter and ventricle masks in the anatomical image resolution. The bottom panel shows correlations of the mean signals calculated from the masks in functional images.

For the alternative processing stream without FastSeg, the correlation was lower than the FastSeg, while they were still higher than 0.9 for most samples with the minimum of 0.89. Although the FastSeg offers a better segmentation quality, the alternative processing stream would have an acceptable quality to make a noise regressor if GPU is not available.

## 5 RTPSpy usage

### 5.1 Building a processing pipeline

#### 5.1.1 Low-level interface

Figure 5 shows a pseudo script to create an online fMRI data processing pipeline. This is presented for explaining the low-level interfaces of building an RTPSpy pipeline. A script with higher-level interfaces using RTP_APP utility methods is presented later (Fig. 6). To make a pipeline, we should create instances of each processing module (Fig. 5A), just set the chaining module in the ‘next_proc’ property (Fig. 5B), and the pipeline is ready. The modules’ combination and the order can be arbitrary, except that RTP_VOLREG must exist before RTP_REGRESS. The head module can be used as an interface to the pipeline (Fig. 5C).

**Figure 5.** A pseudo script of low-level interfaces of pipeline creation with RTPSpy. The comments with bold alphabet indicate a part of the script explained in the main text. To make a pipeline, we should create instances of each processing module (A) and set a chaining module to the ‘next_proc’ property (B). The head module can be used as an interface to the pipeline (C). Properties of the modules in a pipeline can be directly accessed and set by the module instances (D). Calling the ‘redy_proc’ method initializes the pipeline (E). The processing is run by feeding a NiBabel Nifti1Image object to the ‘do_proc’ method of the pipeline (F).

**Figure 6.**
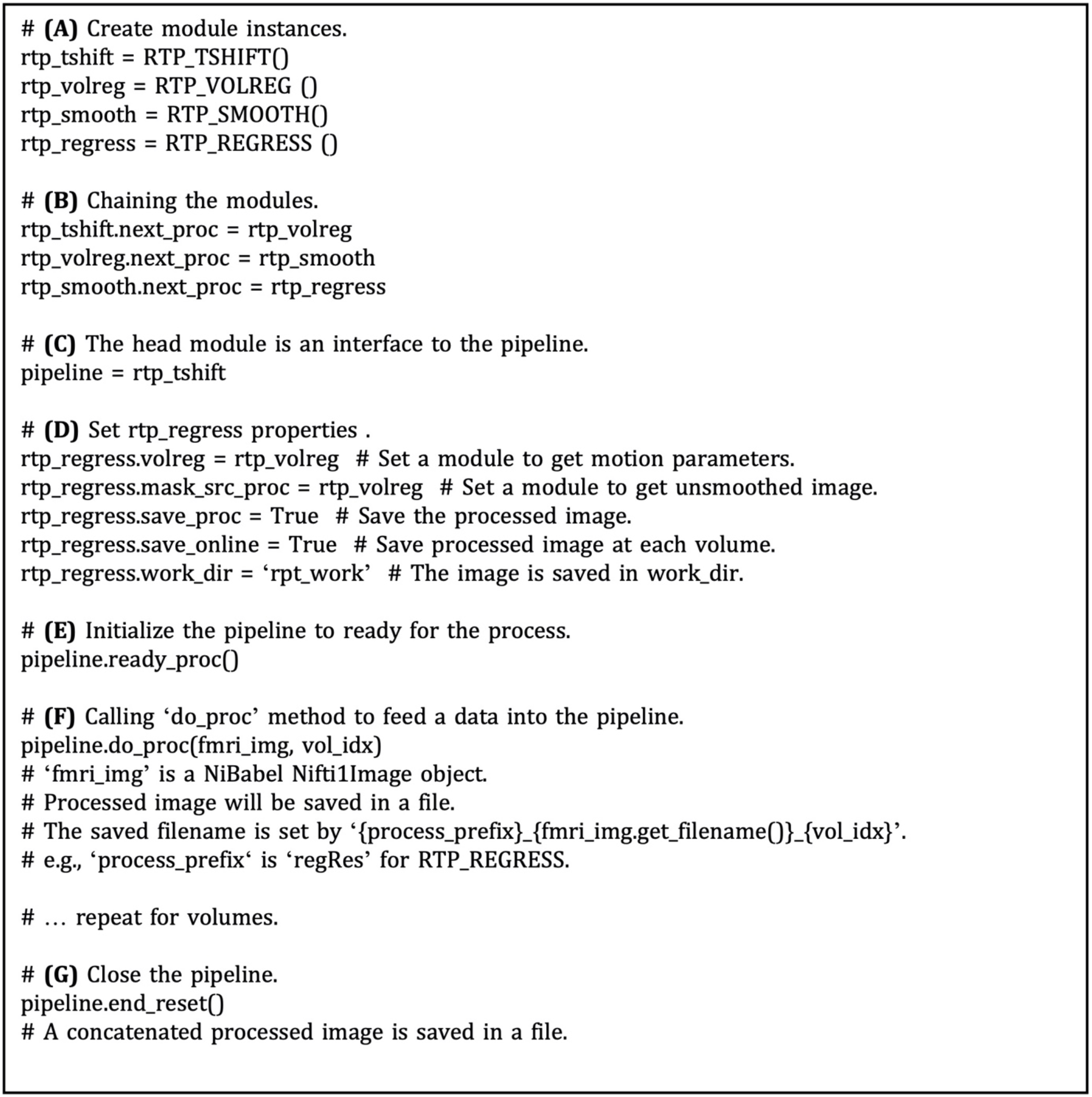
A snippet of an example script to run a real-time processing pipeline using RTP_APP interfaces. The comments with bold alphabet indicate a part of the script explained in the main text. Instantiating the RTP_APP class (A) creates all processing modules inside it. If you use the ‘make_masks’ method to create the mask images, call the method of the rtp_app instance (B). This example uses dummy physiological signals (C). The module properties can be set directly by accessing the ‘rtp_app.rtp_obj’ property (C) or by feeding a property dictionary to the ‘rtp_app.RTP_setup’ method (D and F). If physiological signals are not available, you can disable RETROICOR by setting the ‘phys_reg’ property of ‘REGRESS’ to ‘None’ (E). Calling the ‘rtp_app.ready_to_run’ initializes the pipeline (G). The processing can be started by calling the ‘manual_start’ method (H). To close the pipeline, call the ‘end_run’ method (I).

Properties of the modules in a pipeline can be directly accessed and set by the module instances (Fig. 5D). For example, ‘volreg’ and ‘mask_src_proc’ is set to rtp_regress to receive motion parameters and unsmoothed image data, which are used to create a motion regressor and an average signal regressor in a segmented mask. The ‘save_proc’ property (saving the processed volume data in a NIfTI file) and ‘online_saving’ property (saving is done at each volume) are also set for rtp_regress to save the processed volume image in a file. When the ‘online_saving’ property is set True, the processed image at each volume is saved in real-time. The online saving is done after the downstream processing of the pipeline is completed so as not to affect the real-time performance of the pipeline processing.

Calling the ‘redy_proc()’ method initialize the pipeline (Fig 5E). The processing is run by feeding a NiBabel Nifti1Image object to the ‘do_proc’ method of the pipeline (Fig. 5F). The pipeline is closed by calling the ‘end_reset’ method, and then a concatenated image file is saved in a file. These low-level interfaces could be useful to develop custom input and output modules by users.

#### 5.1.2 High-level interface with RTP_APP

RTPSpy also offers high-level utility methods in RTP_APP. Figure 6 shows a snippet of an example script to run a real-time processing pipeline using RTP_APP interfaces. Refer also to the system check script in the package (rtpspy_system_check.py) for a complete script. Instantiating the RTP_APP class (Fig. 6A) creates all processing modules automatically inside it. The processing modules can be accessed by rtp_app.rtp_obj[‘TSHIFT’] for RTP_TSHIFT, for example. If you use the ‘make_masks’ method to create the mask images, call the method of the rtp_app instance (Fig. 6B). Then, the properties of the mask files, RTP_SMOOTH.mask_file, RTP_REGRESS.mask_file, RTP_REGRESS.GS_mask (global signal mask), RTP_REGRESS.WM_mask, RTP_REGRESS.Vent_mask, and RTP_APP.ROI_mask, are automatically set. The module properties can also be set directly by accessing the ‘rtp_app.rtp_obj’ property (e.g., Fig. 6C) or by feeding a dictionary to the ‘rtp_app.RTP_setup’ method (Fig. 6D and 6F). You can set a custom mask using these interfaces when you want to set a mask without using the ‘make_masks’ method. The order of the pipeline cannot be modified in this interface, but you can disable a specific module by setting the ‘enables’ property False (e.g., Fig. 8A). This example uses dummy physiological signals (Fig. 6C) to simulate the cardiac and respiration signal recordings. If these signals are not available, set the ‘phys_reg’ property of ‘REGRESS’ to ‘None’ (Fig. 6E). All the properties and possible parameters are described in the script files of each module (see Table 2 for the filenames). The pipeline creation is done in the ‘rtp_app. RTP_setup’ method. The ‘save_proc’ property of the last module (e.g., RTP_REGRESS) is automatically set True, and the RTP_APP object is connected after the last module in the ‘rtp_app.RTP_setup’ method.

Calling the ‘rtp_app.ready_to_run’ initializes the pipeline (Fig. 6G). The processing can be started by calling the ‘manual_start’ method (Fig. 6H), and then the RTP_WATCH module starts watching a new file in the watched directory. The start of the processing can also be triggered by a TTL signal implemented in the RTP_SCANONSET module (rtp_scanonset.py file). To close the pipeline, call the ‘end_run’ method (Fig. 6I), then the WATCH module stops monitoring, and the online processed data is saved in a file.

To customize the feedback signal calculation, you can modify the ‘do_proc’ method in RTP_APP (rtp_app.py). The RTP_APP module works as a pipeline terminal, receiving the processed data, extracting the neurofeedback signal, and sending the signal to an external application. By default, it extracts the mean signal in the ROI mask, but it can be overridden according to the individual application need. An example way to make a customized application class is shown in Section 7, Figure 10.

### 5.2 Example rtfMRI session

Figure 7 presents an example procedure of a real-time fMRI (rtfMRI) session. Note, this is not a requirement for the library but only an example of a single-visit session and anatomically-defined neurofeedback target region. The rtfMRI session using RTPSpy could start with an anatomical image scan and a reference functional image scan to make the mask images. We usually perform a restingstate scan after an anatomical scan, and an initial image of the resting-state scan acquired in real-time is used as the reference function image. A pre-acquired anatomical image can also be used in the processing if the study is multi-visits and an anatomical image has been scanned previously. The mask creation using the ‘make_masks’ method in RTP_APP can be finished during the resting-state scan so that no waiting time is required for a participant. If no resting-state scan is necessary, a short functional scan with the same imaging parameters as the neurofeedback runs can also be used. Then, you can set the RTP parameters, run the ‘RTP_setup’ and ‘ready_to_run’ methods, and start the neurofeedback scan.

**Figure 7.**
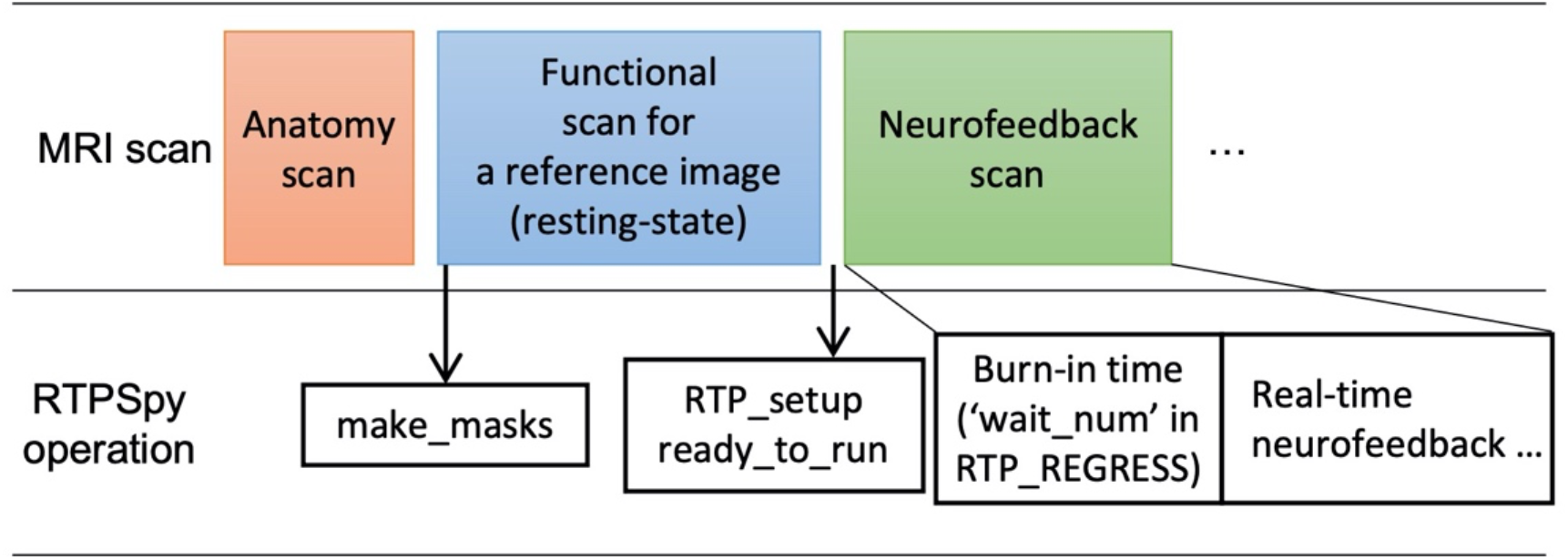
Example real-time fMRI session procedure with RTPSpy. This is an example session and not a requirement for library use.

A critical RTP property related to the task design is the ‘wait_num’ in RTP_REGRESS. This property determines how many volumes the module waits before starting the regression. The task block should start after this burn-in time. Note that this number does not include the initial volumes discarded before the fMRI signal reaches a steady state. The ‘wait_num’ must be larger than the number of regressors, but the just enough number is not enough because the regression with small samples overfits the data, resulting in a very small variance in the denoised output (19). The actual number of required samples depends on the number of regressors and the target region. This waiting time could limit the neurofeedback task design. However, using a noise-contaminated signal as neurofeedback has a high risk of artifactual training effect (6), which may degrade the validity of an experiment. Therefore, the online image processing should include necessary noise regressors, and the task design should accept the initial burn-in time. A simulation analysis would help determine the necessary noise regressors and the optimal number of waiting TRs. Our previous report (7), investigating what brain regions and connectivity were more contaminated with noises as well as the effect of each noise regressor to reduce the noise, may also help. The volumes during the burn-in time are processed at the beginning of RTP_REGRESS processing (thus, the first processing time of RTP_REGRESS could take long, as shown in Figure 4 of Misaki and Bodurka (7)). These volumes can be used, for example, for the baseline calculation to scale the neurofeedback signal (28, 29).

## 6 Simulating real-time fMRI processing

One of the most effective ways to examine the integrity of real-time signal calculation is to simulate online processing and neurofeedback signal calculation using previously obtained fMRI data (7, 15, 16). Assuring the integrity of online noise reduction is critical for neurofeedback training. If the noise reduction is insufficient, other factors than brain activation could confound the training effect (6). Not only for the online image processing, but the feedback signal calculation also can be unique in the online analysis, for example, in the connectivity neurofeedback. The online connectivity calculation should use a short window width for a timely feedback signal reflecting the current brain state, and the optimal window width for the neurofeedback training would be specific to the target region and the task design (15). In addition, simulating the signal processing is useful to evaluate the level of the actual feedback signal. For example, when the baseline level of the neurofeedback signal is adjusted by a mean signal in the preceding rest block (28, 29), simulating such signal calculation could help to estimate a possible signal range to adjust a feedback presentation.

The modular library design of the RTPSpy helps perform a simulation with a simple script. While the simulation can be done by copying the data volume-by-volume into the watched directory, you can also inject the data directly into the pipeline for faster simulation. An example simulation script is provided as the ‘example/Simulation/rtpspy_simulation.py’ file in the package. Figure 8 shows a snippet of the example simulation script. The pipeline creation is the same as shown in Fig. 6 except for disabling the RTP_WATCH module (Fig. 8A) and getting the pipeline object returned from the ‘ready_to_run’ method (Fig. 8B). The simulation can proceed with feeding the Nibabel NiftiImage object to the ‘do_proc’ method of the pipeline (Fig. 8C). This method receives a volume image, image index (optional), and the end time of the previous process (optional). Calling the ‘end_run’ method closes the pipeline and returns the saved filenames (Fig. 8D). The output files include the parameter log (text file), ROI signal time-series (csv file), and the denoise image saved as a NIfTI file. You can modify the neurofeedback signal calculation by overriding the do_proc method in the RTP_APP, as explained in the next section, Figure 10.

**Figure 8.**
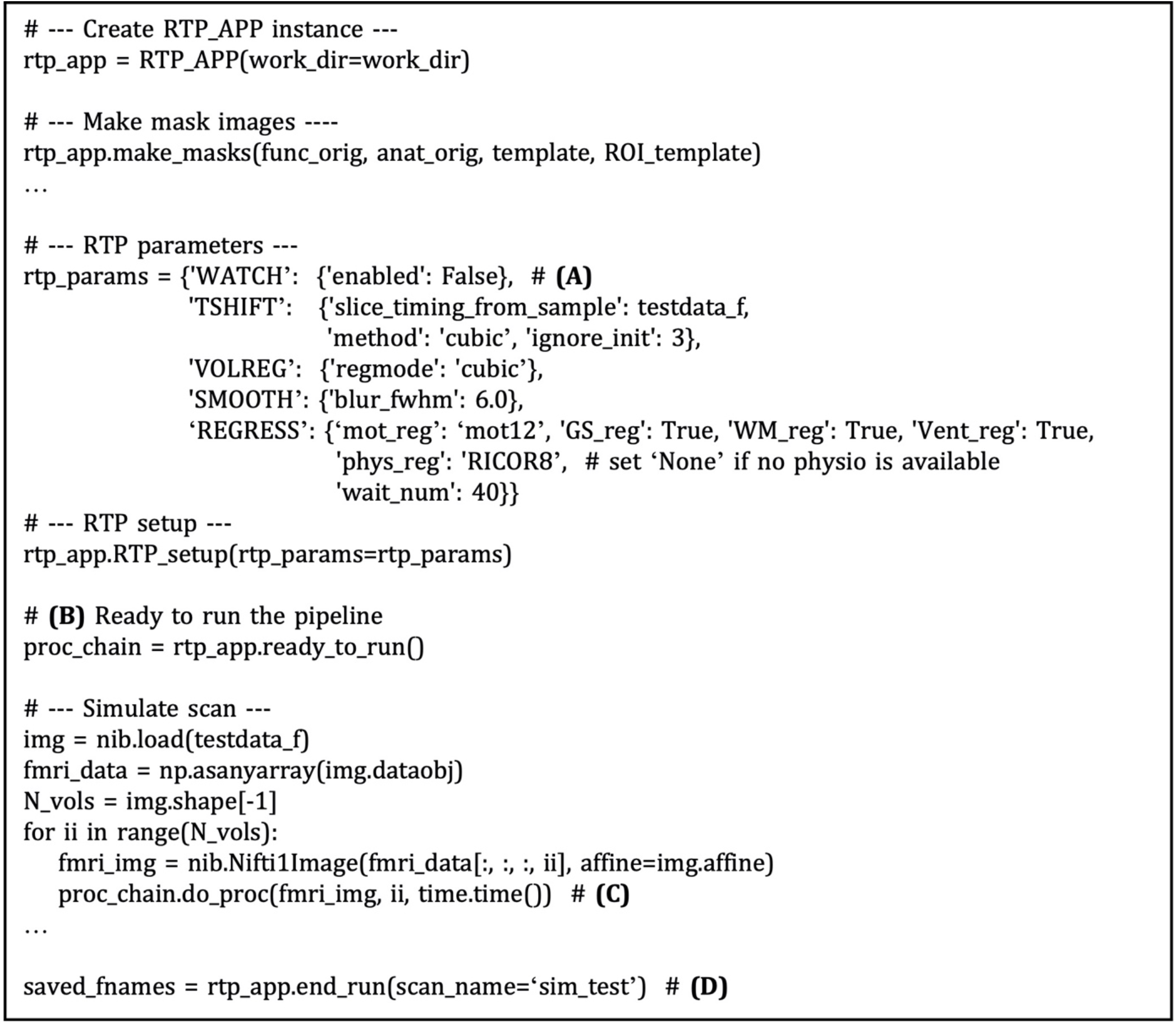
A snippet of an example script of running a real-time fMRI processing simulation with RTPSpy. The comments with bold alphabet indicate a part of the script explained in the main text. The pipeline creation is the same as shown in Fig. 6 except for disabling the RTP_WATCH module (A) and returning the pipeline object from the ‘ready_to_run’ method (B). The simulation can proceed with feeding the Nibabel NiftiImage object to the ‘do_proc’ method of the pipeline (C). Calling the ‘end_run’ method closes the pipeline and returns the saved filenames (D).

## 7 An example GUI application integrating the RTPSpy modules

RTPSpy offers a graphical user interface (GUI) class (RTP_UI, rtp_ui.py) for easy access to the module functions. The example GUI application is provided in the ‘example/ROI-NF’ directory in the package. Figure 9 shows the initial windows of this example application. This application is presented for demonstrating how the RTPSpy library can be used to build a custom application and as a boilerplate for making a custom application by a user. This section explains how these example scripts can be modified to make a custom application. For a step-by-step usage of this application other than scripting, please refer to GitHub; https://github.com/mamisaki/RTPSpy/tree/main/example/ROI-NF.

**Figure 9.**
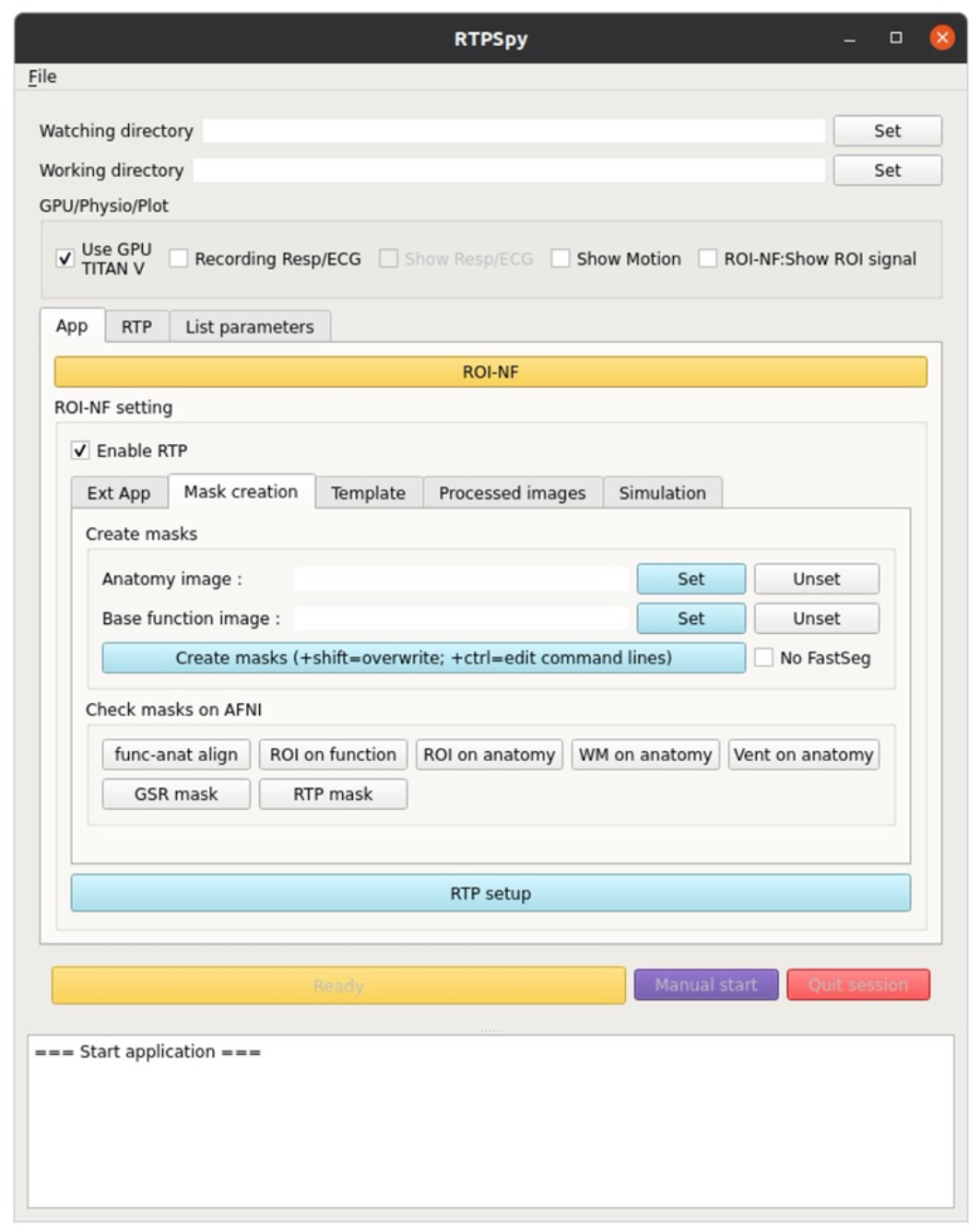
A view of the example GUI application integrating the RTPSpy modules. The figure presents the ‘mask certation’ tab to run the make_masks process with GUI. The example application also offers graphical interfaces to almost all parameters in RTPSpy. Detailed usage of the application is presented in GitHub (https://github.com/mamisaki/RTPSpy/tree/main/example/ROI-NF).

The application development can start with defining a user’s own application class inheriting from the RTP_APP. Figure 10 shows the code snippet from the ‘roi_nf.py’ script file. In this application, ROI_NF class is defined by inheriting RTP_APP class (Fig. 10A). Neurofeedback signal extraction is performed in the ‘do_proc’ method in the ROI_NF class (Fig. 10B). To customize the neurofeedback signal calculation, a user should override this method. The example script calculates the mean value within the ROI mask (Fig. 10D). The ROI mask file is defined in the ‘ROI_orig’ property defined in the RTP_APP class (Fig. 10C). If an external application implements the RTP_SERV module, the signal can be sent to it using the ‘send_extApp’ method (Fig. 10F) by putting a signal value in a specific format string (Fig. 10E).

**Figure 10.**
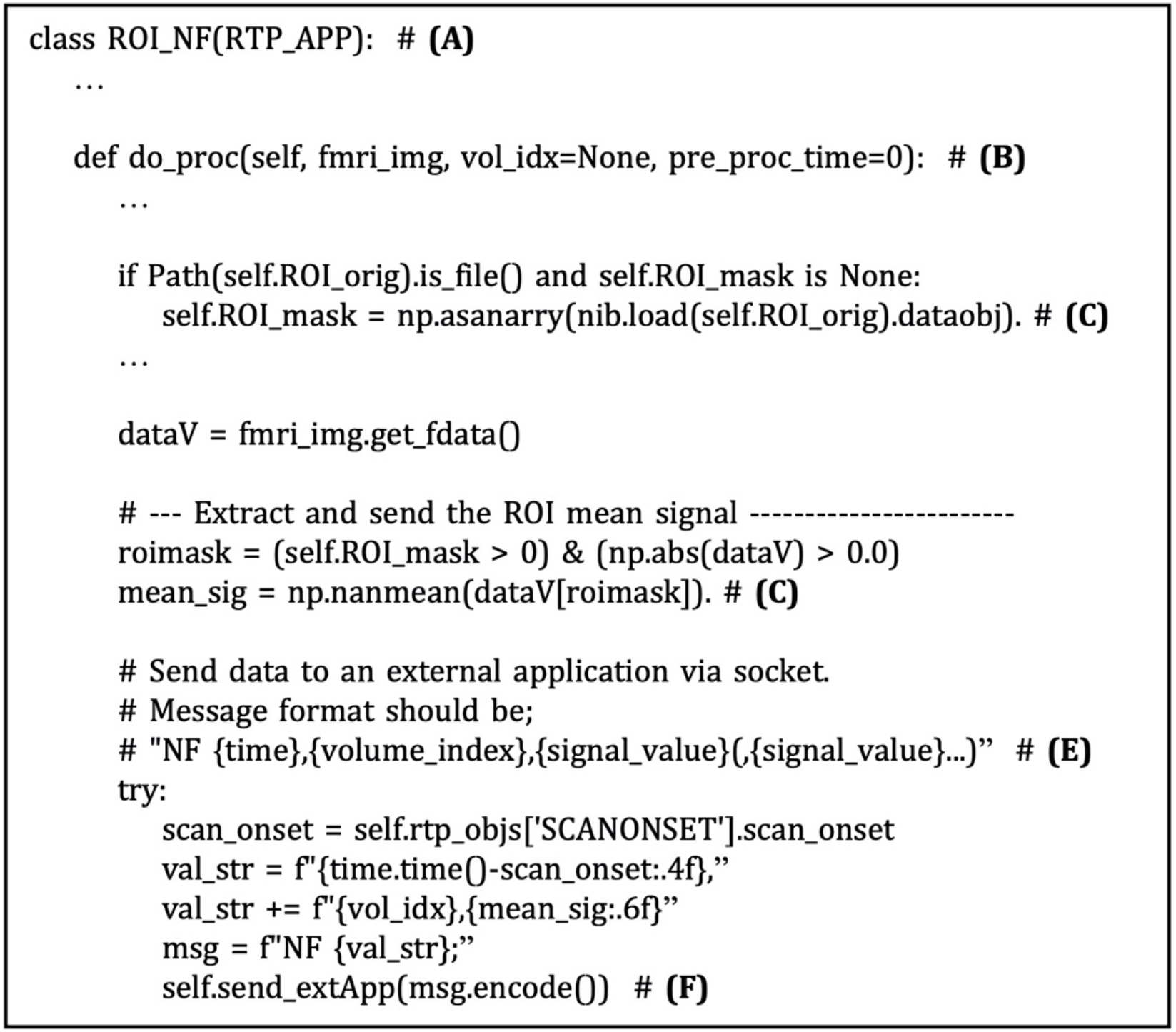
A snippet of an example script of customized neurofeedback signal extraction and sending the signal to an external application in the ‘example/ROI-NF/roi_nf.py’ file. The comments with bold alphabet indicate a part of the script explained in the main text. ROI_NF class is defined by inheriting RTP_APP class (A). Neurofeedback signal extraction is performed in the ‘do_proc’ method (B). The example script calculates the mean value within the ROI mask (D). The ROI mask file is defined in the ‘ROI_orig’ property (C). The signal can be sent to an external application in real-time using the ‘send_extApp’ method (F) by putting it in a specific format string (E).

Figure 11 shows the code snippet of an example external application script for neurofeedback presentation (‘example/ROI-NF/NF_psypy.py’ file). This is an independent PsychoPy (https://www.psychopy.org) application from RTPspy but uses the RTP_SERVE module to communicate with an RTPSpy application. Instantiating the RTP_SERVE class object starts a TCP/IP server running in another thread (Fig. 11A). This class does all the data exchange in the background. The RTP_SERVE object holds the received neurofeedback data in the pandas data frame (https://pandas.pydata.org/)(Fig. 11B). While this example script just displays the latest received value on the screen with text (Fig. 11C), users can modify this part to make a decent feedback presentation.

**Figure 11.**
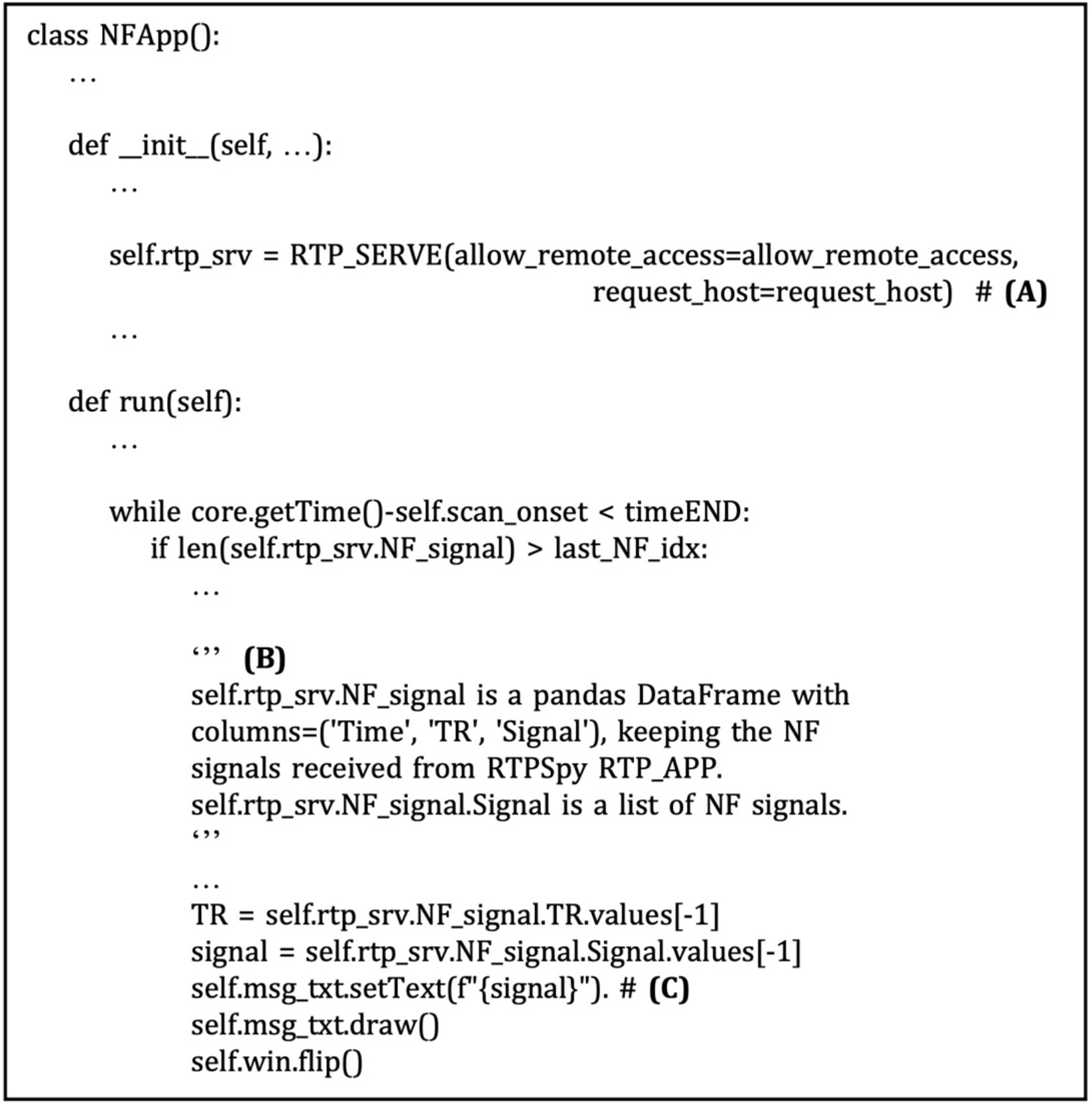
A snippet of an example script of neurofeedback presentation application in the ‘example/ROI-NF/NF_psypy.py’ file. The comments with bold alphabet indicate a part of the script explained in the main text. Instantiating the RTP_SERV class object starts a TCP/IP server running in another thread (A). The RTP_SERVE object holds the received neurofeedback data in the pandas data frame (B). This example script displays the latest received value on the screen with text (C).

In this example application, the GUI operation can be done in parallel to the online image processing as the watchdog in the RTP_WATCH module runs in a separate thread, on which the processing runs. The anatomical image processing tools and a neurofeedback presentation application run on independent processes. Thus, they also run in parallel for a user to operate RTPSpy while an experiment is running. Using these example scripts, a user can develop an easy-to-use and highly customized rtfMRI application with minimum scripting labor. We also provide a full-fledged application of the left-amygdala neurofeedback session (28, 29) in the ‘example/LA-NF’ directory, which is explained in the Supplementary Materials, ‘LA-NF application’ and in GitHub (https://github.com/mamisaki/RTPSpy/tree/main/example/ROI-NF).

## 8 Limitations and environment-specific issues

While the RTPSpy provides general-use libraries for rtfMRI data processing, it is not a complete toolset for all environments. There could be several site-specific settings that a general library cannot support. One of the first critical settings is to obtain a reconstructed MRI image in real-time. The image format, the saved directory, and how to access the data (e.g., network mount or copying to the processing PC) could differ across manufacturers and sites. The RTP_WATCH detects a new file in the watched directory, but setting up the environment to put an fMRI file to an accessible place in real-time is not covered by the library. Specifically, our site uses AFNI’s Dimon command (https://afni.nimh.nih.gov/pub/dist/doc/program_help/Dimon.html) running on the scanner console computer and receives the data sent by Dimon with AFNI’s realtime plugin on a rtfMRI operation computer. This is not a part of the RTPSpy library and may not be possible for all if one cannot install additional software on the scanner console. Users may have to set up real-time access to the reconstructed image according to their environment.

Another caveat of environment-specific implementation is physiological signal recording. One of the advantages of the RTPSpy is its ability to run a physiological noise regression with RETROICOR in real-time. However, the equipment for cardiac and respiration signal recording could vary across the sites and manufacturer. In our site, we measure a cardiac signal using a photoplethysmography with an infrared emitter placed under the pad of a participant’s finger and respiration using a pneumatic respiration belt. These are equipped with the MRI scanner, GE MR750, and the signal is bypassed to the processing PC via serial port. Although we developed an interface class for these signals as RTP_PHYSIO, its implementation is highly optimized for our environment. A user may need to develop a custom script to replace the RTP_PHYSIO module adjusting to the individual environment. Similarly, detection of the TTL signal of a scan start, which is defined in the custom RTP_SCANONSET class, is device-dependent and needs to be implemented by a user according to the user’s device.

RTPSpy depends on several external tools for its anatomical image processing stream. Indeed, the package does not intend to provide an all-around solution by itself. Rather, RTPSpy is supposed to be used as a part of the user’s own application project. A required function specific to each application or environment should be implemented by an external tool or developed by users. We assume that the RTPSpy is used not as a complete package by itself but as a part of a custom application.

## 9 Conclusions

RTPSpy is a library for real-time fMRI processing, including comprehensive online fMRI processing, fast and accurate anatomical image processing, and a simulation system for optimizing neurofeedback signals. RTPSpy focuses on providing the building blocks to make a highly customized rtfMRI system. It also provides an example GUI application wrapped around RTPSpy modules. Although a library package requiring scripting skills may not be easy to use for everyone, we believe that RTPSpy’s modular architecture and easy-to-script interface will benefit developers who want to create customized rtfMRI applications. With its rich toolset and highly modular architecture, RTPSpy must be an attractive choice for developing optimized rtfMRI applications.

## 11 Author Contributions

Masaya Misaki: Conceptualization, Data curation, Formal analysis, Investigation, Methodology, Software, Writing - original draft, Writing - review & editing. Jerzy Bodurka: Conceptualization, Funding acquisition, Investigation, Methodology, Software, Project administration, Supervision. Martin Paulus: Conceptualization, Funding acquisition, Project administration, Supervision, Writing - review & editing.

## 13 Acknowledgments

We dedicate this work in memory and honor of the late Dr. Jerzy Bodurka.

## 14 Data Availability Statement

The RTPSpy package is available from https://github.com/mamisaki/RTPSpy with GPL3 license.

## References

1. Cox RW, Jesmanowicz A, Hyde JS. Real-Time Functional Magnetic Resonance Imaging. Magn Reson Med (1995) 33(2):230–6. doi: 10.1002/mrm.1910330213.

2. Goebel R, Zilverstand A, Sorger B. Real-Time Fmri-Based Brain-Computer Interfacing for Neurofeedback Therapy and Compensation of Lost Motor Functions. Imaging in Medicine (2010) 2(4):407–15.

3. Sulzer J, Haller S, Scharnowski F, Weiskopf N, Birbaumer N, Blefari ML, et al. Real-Time Fmri Neurofeedback: Progress and Challenges. Neuroimage (2013) 76:386–99. Epub 20130327. doi: 10.1016/j.neuroimage.2013.03.033.

4. Mulyana B, Tsuchiyagaito A, Smith J, Misaki M, Kuplicki R, Soleimani G, et al. Online Closed-Loop Real-Time Tes-Fmri for Brain Modulation: Feasibility, Noise/Safety and Pilot Study. bioRxiv (2021):2021.04.10.439268. doi: 10.1101/2021.04.10.439268.

5. Thibault RT, MacPherson A, Lifshitz M, Roth RR, Raz A. Neurofeedback with Fmri: A Critical Systematic Review. Neuroimage (2018) 172:786–807. Epub 20171227. doi: 10.1016/j.neuroimage.2017.12.071.

6. Weiss F, Zamoscik V, Schmidt SNL, Halli P, Kirsch P, Gerchen MF. Just a Very Expensive Breathing Training? Risk of Respiratory Artefacts in Functional Connectivity-Based Real-Time Fmri Neurofeedback. Neuroimage (2020) 210:116580. Epub 20200125. doi: 10.1016/j.neuroimage.2020.116580.

7. Misaki M, Bodurka J. The Impact of Real-Time Fmri Denoising on Online Evaluation of Brain Activity and Functional Connectivity. J Neural Eng (2021) 18(4). Epub 20210701. doi: 10.1088/1741-2552/ac0b33.

8. Goebel R. Brainvoyager--Past, Present, Future. Neuroimage (2012) 62(2):748–56. Epub 20120125. doi: 10.1016/j.neuroimage.2012.01.083.

9. Sato JR, Basilio R, Paiva FF, Garrido GJ, Bramati IE, Bado P, et al. Real-Time Fmri Pattern Decoding and Neurofeedback Using Friend: An Fsl-Integrated Bci Toolbox. PLoS One (2013) 8(12):e81658. Epub 20131202. doi: 10.1371/journal.pone.0081658.

10. Koush Y, Ashburner J, Prilepin E, Sladky R, Zeidman P, Bibikov S, et al. Opennft: An Open-Source Python/Matlab Framework for Real-Time Fmri Neurofeedback Training Based on Activity, Connectivity and Multivariate Pattern Analysis. Neuroimage (2017) 156:489–503. Epub 20170621. doi: 10.1016/j.neuroimage.2017.06.039.

11. Heunis S, Besseling R, Lamerichs R, de Louw A, Breeuwer M, Aldenkamp B, et al. Neu(3)Ca-Rt: A Framework for Real-Time Fmri Analysis. Psychiatry Res Neuroimaging (2018) 282:90–102. Epub 20180928. doi: 10.1016/j.pscychresns.2018.09.008.

12. MacInnes JJ, Adcock RA, Stocco A, Prat CS, Rao RPN, Dickerson KC. Pyneal: Open Source Real-Time Fmri Software. Front Neurosci (2020) 14:900. Epub 20200915. doi: 10.3389/fnins.2020.00900.

13. Kumar M, Michael A, Antony J, Baldassano C, Brooks PP, Cai MB, et al. Brainiak: The Brain Imaging Analysis Kit. https://doiorg/1031219/osfio/db2ev (2020). doi: 10.31219/osf.io/db2ev.

14. Ros T, Enriquez-Geppert S, Zotev V, Young KD, Wood G, Whitfield-Gabrieli S, et al. Consensus on the Reporting and Experimental Design of Clinical and Cognitive-Behavioural Neurofeedback Studies (Cred-Nf Checklist). Brain (2020) 143(6):1674–85. Epub 2020/03/17. doi: 10.1093/brain/awaa009.

15. Misaki M, Tsuchiyagaito A, Al Zoubi O, Paulus M, Bodurka J, Tulsa I. Connectome-Wide Search for Functional Connectivity Locus Associated with Pathological Rumination as a Target for Real-Time Fmri Neurofeedback Intervention. NeuroImage Clinical (2020) 26:102244. Epub 20200312. doi: 10.1016/j.nicl.2020.102244.

16. Ramot M, Gonzalez-Castillo J. A Framework for Offline Evaluation and Optimization of Real-Time Algorithms for Use in Neurofeedback, Demonstrated on an Instantaneous Proxy for Correlations. Neuroimage (2019) 188:322–34. Epub 20181213. doi: 10.1016/j.neuroimage.2018.12.006.

17. Peirce JW. Generating Stimuli for Neuroscience Using Psychopy. Frontiers in neuroinformatics (2008) 2:10. Epub 20090115. doi: 10.3389/neuro.11.010.2008.

18. Henschel L, Conjeti S, Estrada S, Diers K, Fischl B, Reuter M. Fastsurfer - a Fast and Accurate Deep Learning Based Neuroimaging Pipeline. Neuroimage (2020) 219:117012. Epub 20200608. doi: 10.1016/j.neuroimage.2020.117012.

19. Misaki M, Barzigar N, Zotev V, Phillips R, Cheng S, Bodurka J. Real-Time Fmri Processing with Physiological Noise Correction - Comparison with Off-Line Analysis. J Neurosci Methods (2015) 256:117–21. Epub 20150904. doi: 10.1016/j.jneumeth.2015.08.033.

20. Kiebel SJ, Kloppel S, Weiskopf N, Friston KJ. Dynamic Causal Modeling: A Generative Model of Slice Timing in Fmri. Neuroimage (2007) 34(4):1487–96. Epub 2006/12/13. doi: 10.1016/j.neuroimage.2006.10.026.

21. Sladky R, Friston KJ, Trostl J, Cunnington R, Moser E, Windischberger C. Slice-Timing Effects and Their Correction in Functional Mri. Neuroimage (2011) 58(2):588–94. Epub 2011/07/16. doi: 10.1016/j.neuroimage.2011.06.078.

22. Ramot M, Kimmich S, Gonzalez-Castillo J, Roopchansingh V, Popal H, White E, et al. Direct Modulation of Aberrant Brain Network Connectivity through Real-Time Neurofeedback. Elife (2017) 6:e28974. doi: https://doi.org/10.7554/eLife.28974.001.

23. Glover GH, Li TQ, Ress D. Image-Based Method for Retrospective Correction of Physiological Motion Effects in Fmri: Retroicor. Magn Reson Med (2000) 44(1):162–7. Epub 2000/07/14. doi: 10.1002/1522-2594(200007)44:1<162::aid-mrm23>3.0.co;2-e.

24. Bagarinao E, Matsuo K, Nakai T, Sato S. Estimation of General Linear Model Coefficients for Real-Time Application. Neuroimage (2003) 19(2 Pt 1):422–9. Epub 2003/06/20. doi: 10.1016/s1053-8119(03)00081-8.

25. Birn RM, Smith MA, Jones TB, Bandettini PA. The Respiration Response Function: The Temporal Dynamics of Fmri Signal Fluctuations Related to Changes in Respiration. Neuroimage (2008) 40(2):644–54. Epub 20071215. doi: 10.1016/j.neuroimage.2007.11.059.

26. Watanabe T, Sasaki Y, Shibata K, Kawato M. Advances in Fmri Real-Time Neurofeedback. Trends Cogn Sci (2017) 21(12):997–1010. Epub 20171012. doi: 10.1016/j.tics.2017.09.010.

27. Ronneberger O, Fischer P, Brox T. U-Net: Convolutional Networks for Biomedical Image Segmentation. In: Navab N, Hornegger J, Wells WM, Frangi AF, editors. Medical Image Computing and Computer-Assisted Intervention – Miccai 2015. Lecture Notes in Computer Science. Cham: Springer International Publishing (2015). p. 234–41.

28. Young KD, Siegle GJ, Zotev V, Phillips R, Misaki M, Yuan H, et al. Randomized Clinical Trial of Real-Time Fmri Amygdala Neurofeedback for Major Depressive Disorder: Effects on Symptoms and Autobiographical Memory Recall. Am J Psychiatry (2017) 174(8):748–55. Epub 20170414. doi: 10.1176/appi.ajp.2017.16060637.

29. Zotev V, Krueger F, Phillips R, Alvarez RP, Simmons WK, Bellgowan P, et al. Self-Regulation of Amygdala Activation Using Real-Time Fmri Neurofeedback. PLoS ONE (2011) 6(9):e24522.

