## Supplementary materials for "A library for fMRI real-time processing systems in python (RTPSpy) with comprehensive online noise reduction, fast and accurate anatomical image processing, and online processing simulation" (from another version of this preprint: version 1, 2021-12-14)

### *Supplementary Material*

#### 1 ROI-NF application

Example scripts in the 'example/ROI-NF' directory is a GUI application built on the RTPSpy library. This application is provided as a boilerplate of the RTPSpy application to run a neurofeedback session with minimum scripting for custom neurofeedback signal extraction and feedback presentation. Step-by-step instruction on how to use the application is given on the GitHub site; https://github.com/mamisaki/RTPSpy/tree/main/example/ROI-NF#readme.

##### 1.1 Customization of the neurofeedback signal calculation

Neurofeedback signal extraction is performed in the 'do_proc' method in ROI_NF class, defined in the 'roi_nf.py' file. Figure S1 shows the code snippet of signal extraction from a processed fMRI image. To customize the neurofeedback signal calculation, a user should make a new application class inheriting RTP_APP (Fig. S1A) and override the 'do_proc' method (Fig. S1B). The example shown calculates the mean value within the ROI mask (Fig. S1D). The ROI mask file is defined in the 'ROI_orig' property (Fig. S1C), which has been set in the mask creation process (see Section 3 in the main text). The signal can be sent to an external application in real-time using the 'send_extApp' method (Fig. S1F) by putting it in a specific format string (Fig. S1E).

```
class ROI_NF(RTP_APP):   <-- (A)
    …

    def do_proc(self, fmri_img, vol_idx=None, pre_proc_time=0):   <-- (B)
        …

        if Path(self.ROI_orig).is_file() and self.ROI_mask is None:
            self.ROI_mask = np.asanarry(nib.load(self.ROI_orig).dataobj)   <-- (C)
        …

        dataV = fmri_img.get_fdata()

        # --- Extract and send the ROI mean signal -----------------------
        roimask = (self.ROI_mask > 0) & (np.abs(dataV) > 0.0)
        mean_sig = np.nanmean(dataV[roimask])   <-- (D)

        # Send data to an external application via socket.
        # Message format should be;
        # "NF {time},{volume_index},{signal_value}(,{signal_value}...)”   <-- (E)
        try:
            scan_onset = self.rtp_objs['SCANONSET'].scan_onset
            val_str = f"{time.time()-scan_onset:.4f},”
            val_str += f"{vol_idx},{mean_sig:.6f}”
            msg = f"NF {val_str};”
            self.send_extApp(msg.encode())   <-- (F)
```

**Supplementary figure S1.** Code snippet of neurofeedback signal extraction and sending the signal to an external application in example/ROI-NF/roi_nf.py file.

#### 1.2 Customization of the neurofeedback presentation

The neurofeedback presentation application is defined in the 'example/ROI-NF/NF_psypy.py' file. This is an independent PsychoPy (https://www.psychopy.org) application with communication functionality with an RTPSpy application. Figure S2 shows the snippet of the script for the communication. The RTP_SERVE class, defined in the 'rtp_serve.py' file in the RTPSpy package, allows exchanging signals and messages with an RTPSpy application. Instantiating this class object starts a TCP/IP socket server running in another thread (Fig. S2A). This class does all the data exchange in the background. The RTP_SERVE object holds the received neurofeedback data in the pandas data frame (https://pandas.pydata.org/)(Fig. S2B). The example script just displays the latest received value on the screen with text (Fig. S2C). Users can modify this part to make a decent feedback presentation.

```
class NFApp():
    …

    def __init__(self, …):
        …

        self.rtp_srv = RTP_SERVE(allow_remote_access=allow_remote_access,   ← (A)
                                 request_host=request_host)
        …

    def run(self):
        …

        while core.getTime()-self.scan_onset < timeEND:
            if len(self.rtp_srv.NF_signal) > last_NF_idx:
                …

                ‘’’
                self.rtp_srv.NF_signal is a pandas DataFrame with   ← (B)
                columns=('Time', 'TR', 'Signal'), keeping the NF
                signals received from RTPSpy RTP_APP.
                self.rtp_srv.NF_signal.Signal is a list of NF signals.
                ‘’’
                …
                TR = self.rtp_srv.NF_signal.TR.values[-1]
                signal = self.rtp_srv.NF_signal.Signal.values[-1]
                self.msg_txt.setText(f"{signal}")   ← (C)
                self.msg_txt.draw()
                self.win.flip()
```

**Supplementary figure S2.** Code snippet of neurofeedback presentation script in the 'example/ROI-NF/NF_psypy.py' file.

#### 2 LA-NF application

The scrips in the 'example/LA-NF' directory implement the left amygdala neurofeedback session with happy autobiographical memory recall (Zotev et al., 2011;Young et al., 2017). This script is for a demonstration of a full-fledged GUI application using RTPSpy. Note that this application was not used in the previous studies and several parameters are different from the previous reports. Step-by-step instruction on how to use the application is given on the GitHub site; https://github.com/mamisaki/RTPSpy/tree/main/example/LA-NF#readme.

#### 2.1 Task procedures

The neurofeedback run consists of 40s-blocks of Happy, Count, and Rest conditions. At first, the participant is shown the instruction screen (Fig. S3A) that explains each condition. During the Happy condition, the cue 'Happy' and two color bars (red, blue) are displayed on the screen (Fig. S3B). The red bar represents the neurofeedback signal, updated continuously by changing the height based on the neurofeedback signal. The neurofeedback signal is also indicated by a number shown below the red bar. The experiment includes two control conditions ('Count' and 'Rest') to reset the participants' emotional state. During the Count condition (Fig. S3C), a participant is shown the cue 'Count' with the specific instruction to count backward from 300 by subtracting a specified integer. During the Rest condition (Fig. S3D), a participant is presented with the cue 'Rest' and asked to relax while looking at the display screen.

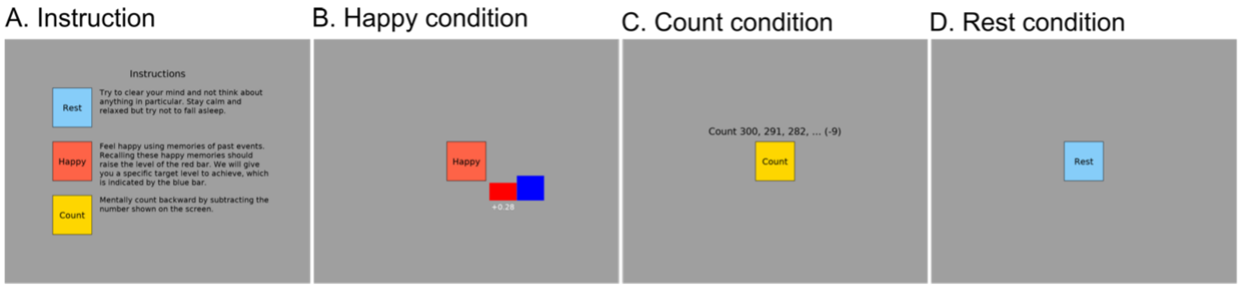


**Supplementary figure S3.** LA-NF task presentation screens.

The neurofeedback signal is the normalized signal subtracting the mean of the preceding Rest period signal and divided by the standard deviation of the Rest period signal. The Rest period signal used for the normalization is the last 24s signals of the 'Rest' block preceding the Happy block. The neurofeedback signal is computed as a moving average of the current and two available preceding values to reduce bar fluctuations. The initial 4s signals in each Happy block are not presented to wait for the hemodynamic response delay. These calculations are implemented in an external application script, 'example/LA-NF/NF_psypy.py'. RTPSPy application only extracts the mean left amygdala signal from the processed image and sends it to the application.

### 3 References

Young, K.D., Siegle, G.J., Zotev, V., Phillips, R., Misaki, M., Yuan, H., Drevets, W.C., and Bodurka, J. (2017). Randomized Clinical Trial of Real-Time fMRI Amygdala Neurofeedback for Major Depressive Disorder: Effects on Symptoms and Autobiographical Memory Recall. *Am J Psychiatry* 174**,** 748-755.

Zotev, V., Krueger, F., Phillips, R., Alvarez, R.P., Simmons, W.K., Bellgowan, P., Drevets, W.C., and Bodurka, J. (2011). Self-Regulation of Amygdala Activation Using Real-Time fMRI Neurofeedback. *PLoS ONE* 6**,** e24522.
